# Identification of leptolin as a novel anti-obesity adipokine

**DOI:** 10.1101/2024.09.03.610963

**Authors:** Jiarui Liu, Bingwei Wang, Zhijie Su, Xiaoxv Han, Miao He, Yun Zhao, Yujia Hou, Daotong Li, Weiguang Zhang, Lihua Qin, Ke Wang, Yanchun Li, Yi Yan, Siwang Yu, Xiaoshuai Huang, Tairan Yuwen, Ruimao Zheng

## Abstract

Adipokines are key factors in regulating energy homeostasis. We identified a novel adipokine; we named it Leptolin. In humans, leptolin levels in white adipose tissue were positively corrected with exercise, and negatively associated with body mass index. Leptolin levels were positively correlated with lipolysis-promoting gene expression. Elevated leptolin in plasma of athletes, whereas lowered leptolin in plasma of obese individuals were observed. Leptolin gene-knockout mice exhibited increased adiposity and body weight, and decreased energy expenditure. Leptolin gene-overexpression mice showed obesity-resistant phenotypes. Treatment with leptolin promoted fat mobilization and energy expenditure, and reduced body weight, without affecting food-intake and motor activity. Together, leptolin, a novel adipokine with a capacity to improve metabolic status, may serve as a new therapeutic agent for obesity and metabolic disorders.

## Main Text

Obesity, a chronic disease defined by excessive fat accumulation, has become an important cause of morbidity and mortality in humans(*1*). Obesity results from the dysregulation of energy balance(*2, 3*). Adipokines, as proteins secreted by adipose tissue, play critical roles in the regulation of energy balance(*4*). For instance, the leptin, an anti-obesity adipokine, activates hypothalamus to promote fat utilization, elevates energy expenditure (EE), and reduces appetite and body weight(*5-7*). The adiponectin stimulates fat mobilization and reduces fat accumulation(*8, 9*). Irisin, an exercise-induced adipokine/myokine, promotes lipolysis, induces white adipose browning, and enhances EE(*10, 11*). Adipokines have potential for treatment of obesity and related metabolic disorders(*12*). Leptin replacement therapy improves body weight, endocrine function and behavior in obese patients with leptin deficiency(*13*). Adiponectin has anti-obesity and insulin-sensitizing effects(*14*). Taken together, these lines of emerging evidence reveal that anti-obesity adipokines may stimulate lipolysis, elevate EE and improve metabolic status.

As the conditions for discovering novel adipokines, physical exercise and cold exposure promote fat-burning(*15, 16*), induce synthesis and secretion of anti-obesity adipokines, such as adiponectin and irisin(*10, 11, 17, 18*). Bruce M. Spiegelman and colleagues identify prosaposin as a novel adipokine by using resource of proteome changes in exercise and cold adaptation(*19*). Francesc Villarroya and colleagues identify CXCL14 as a brown fat-secreted adipokine through analysis of transcriptomic data from brown adipose tissue (BAT) in mice exposed to cold(*20*). These “exerkines” have potential roles in improving metabolic health and treating obesity(*21, 22*). In the present study, by analyzing actively transcribed genes in white adipose tissue (WAT) induced by running, swimming and cold exposure, we identify a novel anti-obesity adipokine, the leptolin, encoded by TMEM52 gene. Leptolin enhances EE and prevents high-fat diet (HFD)-induced obesity, without affecting appetite and food intake. Leptolin may have the potential to treat obesity and related metabolic diseases.

## Results

### Tmem52 is identified as a candidate gene induced by fat-burning conditions

Serving as the conditions for discovering novel adipokines, exercise and cold exposure induce secretion of bioactive proteins from adipose tissue(*19, 20*). Schematic illustration showed the strategy for identifying novel anti-obesity adipokines (**fig. S1, fig. S2A**). Genome-wide transcriptome sequencing analysis showed that expression of genes associated with lipolysis (fatp2, fabp3, pdk4 and plin5) and mitochondrial function (pgc1a, cox7a, cox8b and ucp1) were upregulated; and expression of genes involved in lipid synthesis (cebpa and srebf1) and energy metabolism negative regulation (acvr2b, npr3, nr1d1 and tshr) were downregulated in the inguinal white adipose tissue (iWAT) under these conditions (**fig. S2B**). Intriguingly, gene expression of the leptin and the asprosin was decreased, and gene expression of the irisin was increased in iWAT under these fat-burning conditions (**fig. S2B**). Leptin, asprosin and irisin have previously been reported as being secreted from white adipose tissue under the influence of cold exposure and exercise(*10, 11, 23-26*). Venn analysis showed that 424 genes were upregulated and 295 genes were downregulated (**fig. S2C**). Gene ontology (GO) and Kyoto encyclopedia of genes and genomes (KEGG) analyses showed that the pathways associated with lipolysis, β-oxidation and mitochondrial function were activated (**fig. S2D**). Gene set enrichment analysis (GSEA) demonstrated that exercise and cold exposure promoted the lipid and glucose metabolism processes (**fig. S2E**). Tmem52, a gene of unknown function, was among the selected top 20 upregulated genes in response to exercise and cold exposure (**Fig. 1A-B**). Notably, genetic correlation analysis between obesity and over 16000 genes showed that tmem52 was as one of the top-ranking genes (**Fig. 1C**). Signal peptide prediction showed that the short N-terminal amino acid sequences of human and mouse tmem52-encoded protein were highly likely to be a signal peptide (**Fig. 1C**). Importantly, tmem52 gene expression was positively associated with the genes related to mitochondrial function and fat mobilization, including cox7α1, cox8β, pgc1α, ucp1 and fatp2 (**fig. S2G**). Real-time quantitative PCR analysis validated that tmem52 and the genes associated with mitochondrial function (pgc1α, cox8β, cox7α1, ucp1, prdm16) and fat mobilization (plin1, fatp1, fatp2) were upregulated in response to exercise and cold exposure (**fig. S2H**). Together, these findings identify tmem52 as a candidate gene in white adipose tissue under fat-burning conditions.

**Fig. 1.**
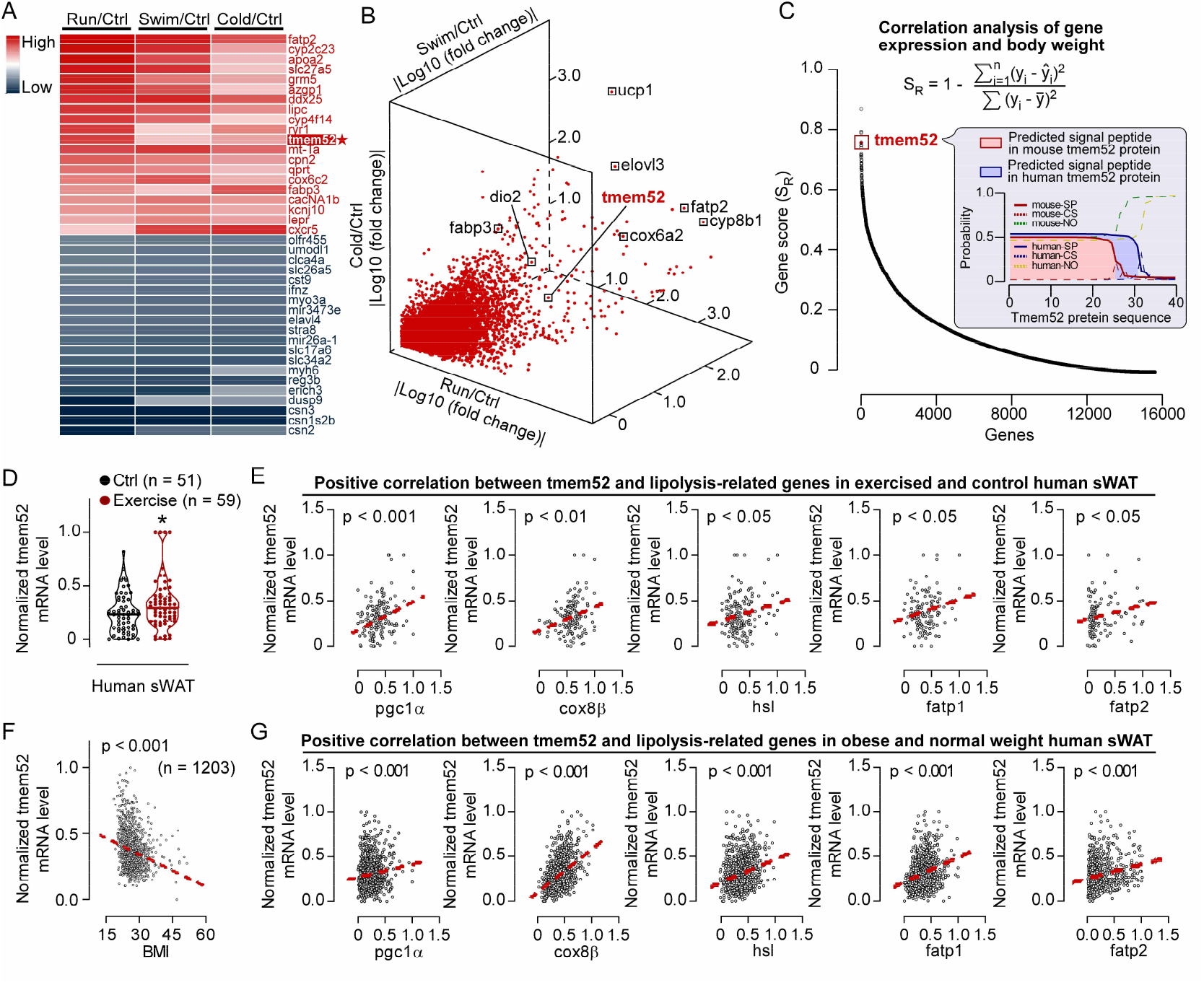
Tmem52 is identified as a candidate gene induced by fat-burning conditions. **(A-C)** RNA sequencing of iWAT of mice in response to exercise and cold exposure; Ctrl, n = 3, Run, n = 3, Swim, n = 3, Cold, n = 3. **(A)** Heatmaps representing the selected 20 upregulated (red) and 20 downregulated (blue) genes -run versus ctrl group mice (left), swim versus ctrl group mice (middle) and cold versus ctrl group mice (right). **(B)** Three-dimensional plot highlighting the differential expression genes (DEGs). **(C)** Distribution of the calculated score (correlation between gene expression and body weight) of individual genes, and signal peptide prediction based on the amino acid sequence of the human and mouse tmem52-encoded protein. The red dot represented tmem52 gene. SP, signal peptide, CS, cleavage site, NO, non-signal peptide or cleavage site. **(D-G)** Analysis of the human sWAT datasets. **(D)** Tmem52 gene expression in sWAT of differently normalized Ctrl/Exercise human datasets (Ctrl, n = 51, Exercise, n = 59); Student’s t-test was used for statistical analysis; *p < 0.05. **(E)** Scatter plots of mRNA levels of tmem52 and genes associated with mitochondrial function and fat mobilization in sWAT of differently normalized Ctrl/Exercise human datasets; The red line showed correlation. **(F)** Scatter plots of BMI and tmem52 mRNA levels in sWAT of differently normalized large sample human datasets (n = 1203); The red line showed correlation. **(G)** Scatter plots of mRNA levels of tmem52 and lipolysis-related genes in sWAT of differently normalized large sample human datasets; The red line showed correlation.

### Tmem52 mRNA levels in subcutaneous WAT are increased in athletes, and lowered in obese individuals

To explore the relevance of tmem52 to human obesity, we analyzed the whole-genome sequencing data of human subcutaneous WAT (sWAT). Schematic diagram showed the enrollment criteria for exercised human sWAT transcriptome datasets (**fig. S3A**). Elevated tmem52 gene expression levels in sWAT of humans with exercise were observed, as compared with that of non-exercised humans (**Fig. 1D**). Intriguingly, tmem52 mRNA levels were positively correlated with the gene expressions involved in mitochondrial function and lipolysis (pgc1α, cox8β, hsl, fatp1, fatp2) in sWAT of these non-exercised and exercised humans (**Fig. 1E**). A negative correlation between tmem52 mRNA levels in sWAT and body mass index (BMI) was found in combined two large-sample datasets of normal weight and obese humans (**fig. S3B, Fig. 1F**). Likewise, tmem52 mRNA levels were also positively correlated with the lipolysis-promoting genes (pgc1α, cox8β, hsl, fatp1 and fatp2) in sWAT of these humans (**Fig. 1G, fig. S4A**). Further, a negative correlation between tmem52 and leptin at mRNA level was detected in sWAT of these normal weight and obese humans (**fig. S4B**). Together, these findings uncover an increase in tmem52 mRNA level in WAT of athletes as well as a decrease in tmem52 mRNA level in sWAT of obese individuals, suggesting a therapeutic potential of tmem52 for obesity treatment in humans.

### Leptolin (tmem52-encoded protein) is present in human and mouse plasma

We then evaluated the evolution of tmem52. Unequivocal homologs of tmem52 were identified in mammals (**fig. S5A**). Sequence alignment analysis showed a conserved tmem52 amino acid sequence among mammal species (**fig. S5B**). Human and mouse tmem52-encoded protein are 77% identical (**fig. S5C**), compared to 100% identity for irisin, 83% identity for leptin, 83% identity for adiponectin and 59% identity for resistin, suggesting a conserved function of tmem52 in human and mouse(*10*).

Considering that both human and mouse tmem52 mRNA levels in WAT were inversely correlated with obesity, we named the tmem52 gene-encoded protein as leptolin. An immunogenic antigen was selected for the generation of a functional and highly specific anti-leptolin antibody (**fig. S6**). With the predicted signal peptide structure of leptolin in mind (**Fig. 1C**), we detected the leptolin in plasma of humans and mice (**Fig. 2A-C**). In adult athletes undergoing supervised endurance exercise training, a 38.4% increase in the circulating leptolin levels was observed compared with the non-athletes (**Fig. 2A**). In obese (BMI ≥ 28) individuals, a reduction of about 33% in leptolin levels was detected in plasma compared with the normal weight individuals (**Fig. 2B**). Interestingly, a negative correlation between circulating leptolin levels and BMI in humans was found (**Fig. 2B**). In mice, we observed elevated (about 2-fold) plasma concentrations of leptolin under the influence of exercise and cold exposure (**Fig. 2C**). Taken together, leptolin is a secretory protein, and it may have anti-obesity function in humans and mice.

**Fig. 2.**
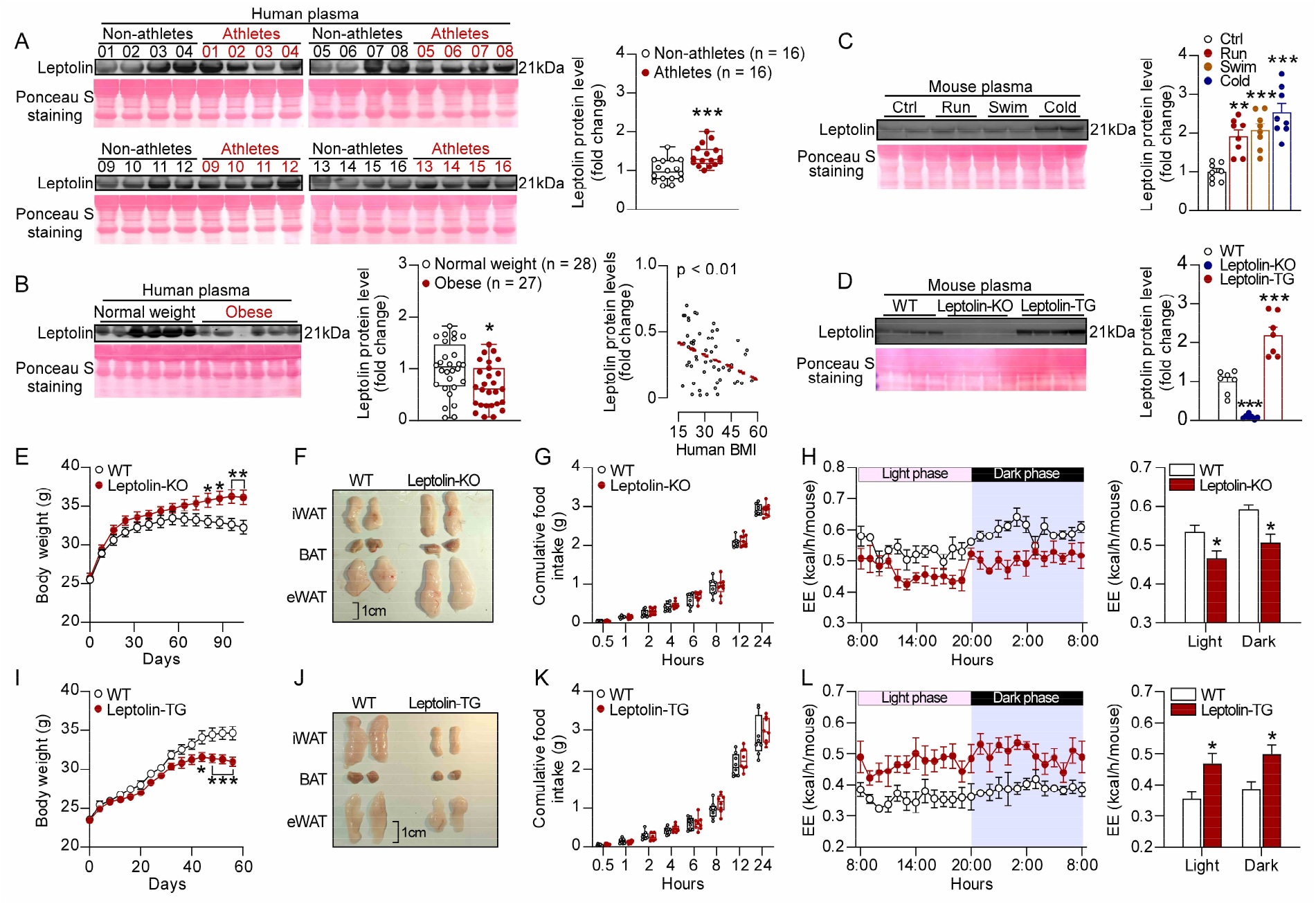
Leptolin deficiency increases the susceptibility to HFD-induced obesity, while leptolin gene overexpression prevents HFD-induced obesity. **(A)** Representative immunoblots of leptolin and images of ponceau s staining from plasma of human athletes and non-athletes; n = 16 per group. **(B)** Representative immunoblots of leptolin and images of ponceau s staining from plasma of normal weight and obese humans, and the scatter plots of BMI and leptolin protein levels in human plasma; normal weight, n = 28, obese, n = 27. **(C)** Representative immunoblots of leptolin and images of ponceau s staining from plasma of mice under fat-burning conditions; Ctrl, n = 8, Run, n = 8, Swim, n = 8, Cold, n = 8. **(D)** Representative immunoblots of leptolin and images of ponceau s staining from plasma of WT, leptolin-KO and leptolin-TG mice; n = 7 per group. **(E-H)** 8-week-old leptolin^+/+^ mice (WT) and leptolin^−/−^ mice (leptolin-KO) were fed a high-fat-diet, tissue harvest was performed at month 3; WT, n = 8, Leptolin-KO, n = 9. **(E)** Body weight. **(F)** Representative images of fat pads (iWAT, eWAT and BAT). **(G)** Cumulative food intake. **(H)** Indirect calorimetry was performed to quantify EE during complete 24 hr light-dark cycles; n = 5 per group. **(I-L)** 8-week-old wild type mice (WT) and leptolin gene overexpression mice (leptolin-TG) were fed a high-fat-diet, tissue harvest was performed at month 2; WT, n = 8, Leptolin-TG, n = 9. **(I)** Body weight. **(J)** Representative images of fat pads (iWAT, eWAT and BAT). **(K)** Cumulative food intake. **(L)** Indirect calorimetry was performed to quantify EE during complete 24 hr light-dark cycles; n = 5 per group. Data were mean ± s.e.m.; Statistical analysis: one-way ANOVA with Bonferroni test for **(C)** and **(D)**, and student’s t test for the rest; *p < 0.05, **p < 0.01, ***p < 0.001.

### Leptolin deficiency increases the susceptibility to HFD-induced obesity, while leptolin gene overexpression prevents HFD-induced obesity

To explore whether leptolin may reduce adiposity and promote energy expenditure, we designed and created the leptolin-deficient (leptolin gene knockout, leptolin-KO) mice (**fig. S7A-B**) and the leptolin gene overexpression (leptolin transgenetic, leptolin-TG) mice (**fig. S7C-D**). Leptolin protein levels in plasma were approximately doubled in leptolin-TG mice compared with wild-type mice, and the leptolin was not detected in plasma of leptolin-KO mice (**Fig. 2D**).

We found that the body weight of leptolin-KO mice was higher than that of controls, equivalent to a total weight gain of 11.9% at month 3 (**Fig. 2E**). The fat-pad weight of iWAT and epididymal WAT (eWAT) was increased in leptolin-KO mice, while the weight of brown adipose tissue (BAT) was not altered (**Fig. 2F, fig. S8A-B**). The size of adipocytes in iWAT of leptolin-KO mice showed about 27.3% larger than that of controls (**fig. S8C**). The cumulative food intake did not differ between groups (**Fig. 2G**). A reduced whole-body EE (**Fig. 2H**) and an increased respiratory exchange ratio (RER) (**fig. S8D**) were observed in leptolin-KO mice, but there was no change in the daily locomotor activity (**fig. S8E**). The gene expression associated with mitochondrial function (pgc1α, cox8β, cox7α1, ucp1, and prdm16) and lipolysis (plin1, fatp1, fatp2, hsl, and atgl) was downregulated in iWAT of leptolin-KO mice (**fig. S8F**). Protein levels of Pgc1α, a key regulator for mitochondrial biogenesis, and p-HSL, a canonical marker of lipolysis, were decreased in iWAT as compared with controls (**fig. S8G**). After 90 days on the high-fat diet, the blood glucose levels of leptolin-KO mice were higher than that of control mice during intraperitoneal glucose tolerance test (GTT) (**fig. S9A-C**); the insulin, triglyceride and total cholesterol levels in plasma of leptolin-KO mice were higher than that of control mice (**fig. S9D, fig. S10A-C**). At thermoneutrality, the body weight and fat mass of leptolin-KO mice were also raised, while the lean mass was not affected (**fig. S11A-D**). Together, leptolin deficiency reduces EE and increases the susceptibility to high-fat diet-induced obesity, suggesting a remarkable energy expenditure-raising function of leptolin.

Conversely, leptolin gene overexpression effectively prevented the HFD-induced obesity with a 10.6% reduction in body weight, as compared with controls at month 2 (**Fig. 2I**). The fat-pad weight of iWAT and eWAT was reduced (**Fig. 2J, fig. S12A-B**). The size of adipocytes in iWAT of leptolin-TG mice was about 11.8% smaller than that of controls (**fig. S12C**). The food intake was not altered (**Fig. 2K**). Notably, leptolin gene overexpression led to an increased EE and a decreased RER (**Fig. 2L, fig. S12D**); and the locomotor activity was not affected (**fig. S12E**). The mRNA levels of the genes involved in mitochondrial function (pgc1α, cox8β, cox7α1, ucp1, and prdm16) and lipolysis (plin1, fatp1, and fatp2), and the protein levels of Pgc1α and p-HSL were enhanced in iWAT of leptolin-TG mice (**fig. S12F-G**). After 60 days on the HFD, leptolin-TG mice exhibited a lowered blood glucose level during GTT and insulin tolerance test (ITT) (**fig. S13A-C**). The plasma insulin and triglyceride levels were reduced (**fig. S13D, fig. S14A-B**); the total cholesterol level in plasma was not changed (**fig. S14C**). Taken together, these observations demonstrate that leptolin gene overexpression elevates EE prevents the development of HFD-induced obesity, showing a therapeutic potential of leptolin in obesity and related metabolic disorders.

### Leptolin is identified as an adipokine

Real-time quantitative PCR analysis showed that the tmem52 mRNA levels in fat pads, especially in iWAT, were elevated under the influence of exercise and cold exposure (**Fig. 3A**). By using the RNAscope assay, a novel RNA in situ hybridization technique, we observed tmem52 mRNA signals in adipocytes of iWAT with high sensitivity and specificity (**fig. S15, Fig. 3B**). The tmem52 mRNA signals in adipocytes were robustly enhanced under the fat-burning conditions (**Fig. 3B, fig. S16A**). Western blot and immunofluorescence analysis validated that the leptolin protein levels in iWAT were elevated under the fat-burning conditions (**fig. S16B-C**).

**Fig. 3.**
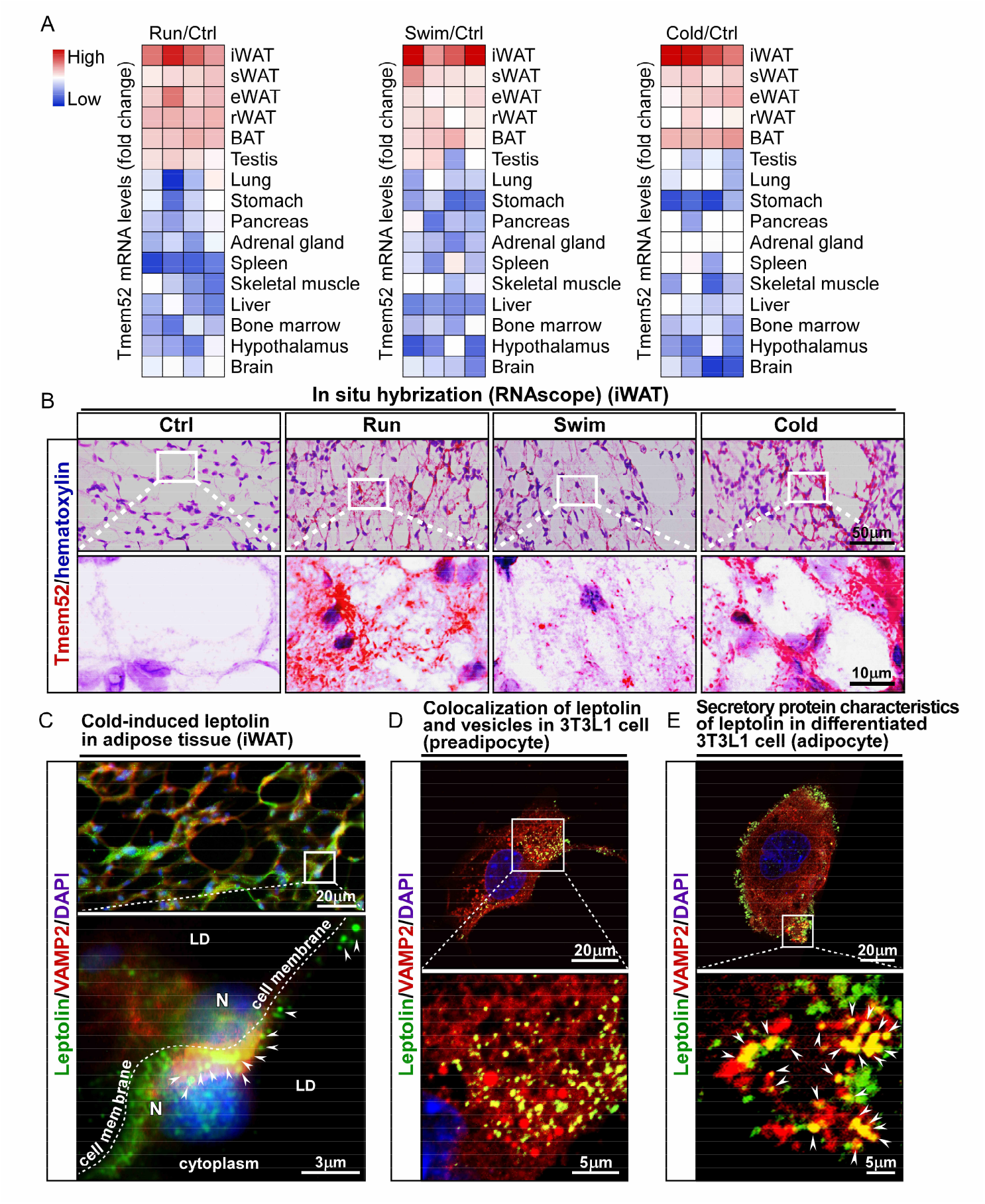
Leptolin is identified as an adipokine. **(A)** Relative mRNA expression (fold change) of tmem52 in multiple organs and tissues of mice in response to exercise and cold exposure; Ctrl, n = 4, Run, n = 4, Swim, n = 4, Cold, n = 4. **(B)** Representative RNAscope images visualizing tmem52 mRNA in iWAT; Red dots were indicative of tmem52 signal with each dot corresponding to a single RNA molecule; Hematoxylin was used to stain nucleus; Scale bar, 50 μm (upper panel) and 10 μm (down panel). **(C)** Representative immunofluorescence images of leptolin and VAMP2 in iWAT; Scale bar, 20 μm (upper panel) and 3 μm (down panel). **(D-E)** Representative immunofluorescence images of leptolin and VAMP2 in undifferentiated 3T3L1 cell **(D)** and differentiated 3T3L1 cell **(E)**; Scale bar, 20 μm (upper panel) and 5 μm (down panel).

Further, immunofluorescence technique visualized leptolin around nucleus and cell membrane in white adipocytes of iWAT (**Fig. 3C**) and in 3T3L1 cells (**Fig. 3D-E**). The co-localization of leptolin and vesicle-associated membrane protein 2 (VAMP2) in iWAT and 3T3L1 cells showed that synthesized leptolin might be transported by secretory vesicles in adipocytes (**Fig. 3C-E**). The distribution of leptolin protein nearby the cell membrane of differentiated 3T3L1 adipocytes mirrored the morphological feature of secreted protein (**Fig. 3E**). Likewise, SR fluorescence-assisted diffraction computational tomography (SR-FACT)(*27*) also observed the leptolin-EGFP signals in vesicles of live 293T cells transfected with plasmid (**fig. S17, movie. S1**). Taken together, leptolin has the typical characteristics of secretory protein in adipocytes. Therefore, we identify leptolin as an adipokine.

### Leptolin administration enhances EE and prevents HFD-induced obesity

Meta-analysis of the 16 obesity-related human studies showed that tmem52 mRNA levels in sWAT of obese individuals were lowered, as compared with that of normal weight individuals (P < 0.00001) (**Fig. 4A-B**). This result strongly suggests leptolin as promising anti-obesity adipokine.

**Fig. 4.**
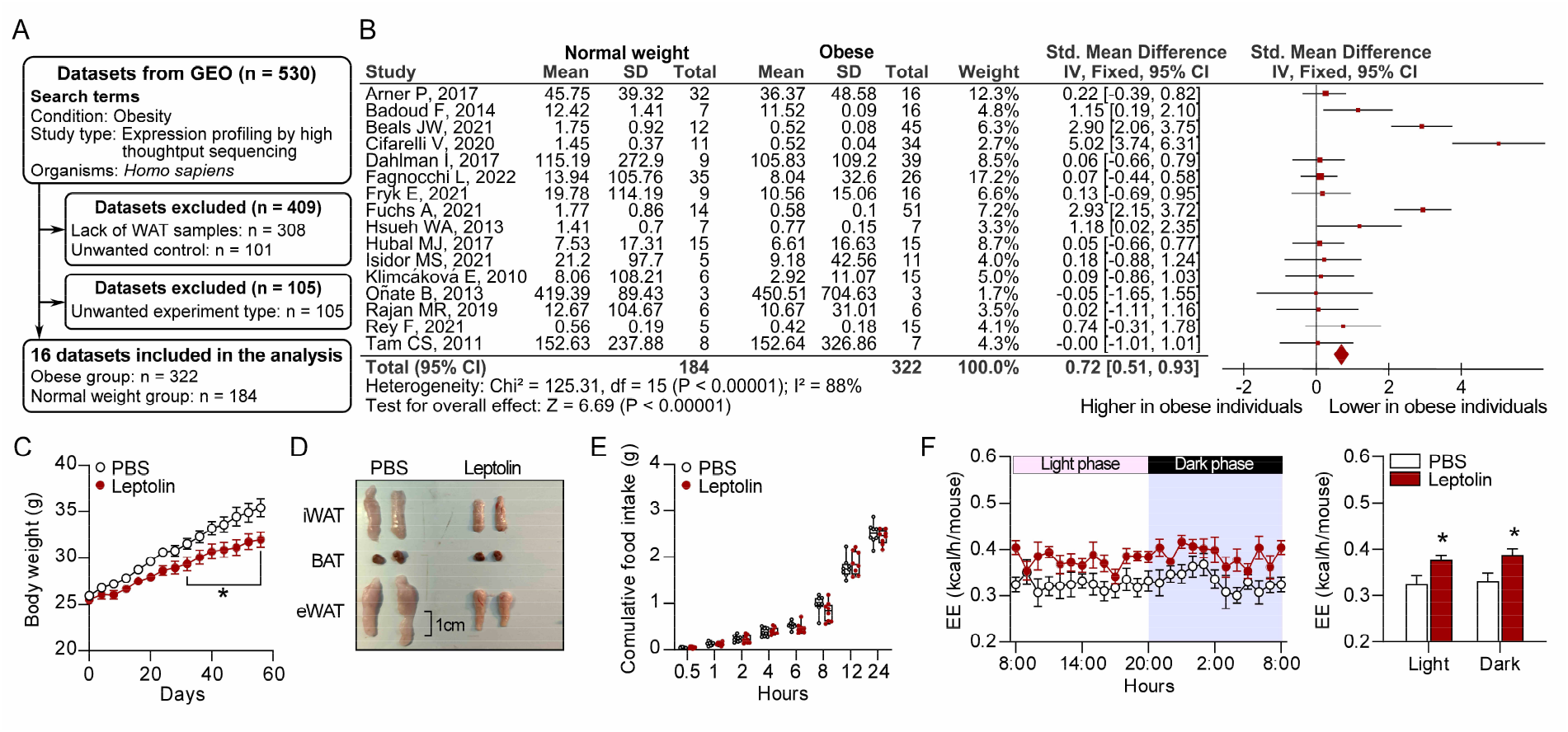
Leptolin administration enhances EE and prevents HFD-induced obesity. **(A)** Schematic illustration of screening workflow of Normal weight/Obese human sWAT datasets, 16 datasets are included in the meta-analysis. **(B)** Forest plot of tmem52 mRNA levels in sWAT of Normal weight/Obese human datasets (Normal weight, n = 184, Obese, n = 322). **(C-F)** 8-week-old wild-type mice were fed a high-fat-diet, received PBS for 4 days as acclimation, then were intraperitoneally injected with leptolin or PBS daily for 2 months, tissue harvest was performed at month 2; PBS, n = 8, Leptolin, n = 8. **(C)** Body weight. **(D)** Representative images of fat pads (iWAT, eWAT and BAT). **(E)** Cumulative food intake. **(F)** Indirect calorimetry was performed to quantify EE of PBS and leptolin-treated mice during complete 24 hr light-dark cycles; n = 6 per group. Data were mean ± s.e.m.; Student’s t-test was used for statistical analysis; *p < 0.05.

To evaluate the therapeutic potential of leptolin in treating obesity, we cloned and produced a leptolin protein fragment (68-196aa) for intraperitoneal (I.P.) injection (**fig. S18A**). The dose-effect study showed that leptolin injection at the dosage levels of 0.5 or 5 mg/kg/d for 2 months effectively reduced body weight and elevated EE in HFD-fed mice, without altering daily food intake (**fig. S18B-E**). The leptolin-treated mice did not display adverse reaction, and there was no apparent toxicity in major organ systems. Thus, leptolin I.P. injection (0.5 mg/kg/d, once daily for 2 months) was selected for study of its anti-obesity effects (**fig. S19A**). We found that leptolin treatment markedly reduced body weight in HFD-fed mice, equivalent to a total weight loss of 10.8% at month 2 (**Fig. 4C**). Leptolin treatment led to a reduced fat-pad weight of iWAT and eWAT (**Fig. 4D, fig. S19B**), and a smaller (about 20.5%) size of adipocytes in iWAT (**fig. S19C**). Cumulative food intake was not altered by leptolin injection (**Fig. 4E**). Leptolin enhanced EE and decreased RER, and did not change the motor activity (**Fig. 4F, fig. S19D-E**). Increased gene expression associated with mitochondrial function (pgc1α, cox8β, cox7α1, ucp1, and prdm16) and lipolysis (plin1, fatp1, and fatp2), and elevated protein levels of Pgc1α and p-HSL were observed in iWAT of leptolin-treated mice (**Fig. 19F-G**). After 60 days of leptolin treatment, an improved blood glucose levels during GTT and ITT were detected (**fig. S20A-C**). The plasma insulin and triglyceride levels were lowered (**fig. S20D, fig. S21A-B**). The plasma total cholesterol level did not differ between groups (**fig. S21C**). Together, leptolin administration enhances EE and reduces weight gain, uncovering that leptolin may hold promise in obesity treatment. Exploring the clinical utility of leptolin may shed new light on the therapy for obesity and related disorders.

## Discussion

We detected the existence of leptolin protein in blood plasma of humans and mice, characterizing leptolin as a secreted protein(*10, 28*). We observed that leptolin levels in WATs, but not in other tissues or organs, were elevated under the conditions of physical exercise and cold exposure. The RNAscope technology, a novel in situ hybridization assay(*29*), revealed that leptolin was an adipocyte-derived hormone. Leptolin protein could be synthesized around nucleus and transported by secretory vesicles in adipocytes. Taken together, we identified leptolin as an adipokine.

As an adipokine, leptolin could be produced by adipocytes and secreted into plasma in mice. In humans, leptolin also could be detected in sWAT and blood plasma. Meta-analysis of global collection of human data showed that leptolin mRNA levels were lowered in sWAT of obese individuals, and elevated in sWAT of athletes. Of note, our experimental data showed decreased levels of leptolin in blood plasma of obese individuals, and raised levels of leptolin in blood plasma of athletes. Human leptolin levels in sWAT and plasma were negatively associated with BMI, which is similar to the feature of other anti-obesity adipokines such as adiponectin(*30, 31*). These findings uncover that leptolin may be a potential anti-obesity adipokine, and it may have the capacity to improve metabolic status.

Leptolin promoted fat utilization, elevated EE and prevented obesity. We observed that leptolin gene-overexpression mice showed elevated lipolysis, enhanced EE, and obesity-resistant phenotypes. Conversely, leptolin gene-knockout mice exhibited adverse phenotypes. Administration of exogenous leptolin promoted fat mobilization, increased EE and led to a total weight loss of 10.8% at month 2 in HFD-induced obese mice. Stimulation of fat utilization and elevation of EE are important therapeutic strategies for obesity(*3, 32, 33*). The leptin, a canonical adipokine, can promote lipolysis and increase basal metabolic rate (BMR)(*5, 34*). Irisin, an adipokine/myokine, can act on adipocytes to trigger lipolysis and elevate EE(*10, 35, 36*). These findings suggest that enhancement of lipolysis and catabolic metabolism and the elevation of EE may underlie the anti-obesity effects of leptolin.

Adipokines are crucial in the regulation of lipid catabolism(*37*). Dysfunction of adipokines contributes to obesity and related complications(*37*). Adipokines are potential therapeutic agents in obesity(*38*). For instance, leptin deficiency results in severe obesity(*39*); Leptin replace therapy promotes lipolysis, elevates EE, and inhibits food intake to reduce body weight(*5, 13*). Adiponectin levels in plasma are lowered in obesity(*31*); Adiponectin treatment enhances EE, reduces food intake and decreases body weight(*8, 9*). Administration of irisin activates white adipocytes and elevates EE to reduce body weight(*10*). In this study, we detected a decreased level of leptolin in blood plasma of obese patients. Moreover, leptolin deficiency augmented fat-pad weights and body weight. Notably, leptolin administration stimulated lipolysis, enhanced EE and reduced body weight. Collectively, we suggest that leptolin may be a novel agent for treating obesity.

Obesity is primarily the result of dysregulation of energy intake and energy expenditure(*3, 40*). To date, the anti-obesity drugs are usually associated with appetite suppression(*41, 42*). Semaglutide mimics the hormone glucagon-like peptide 1 (GLP-1) to inhibit appetite and reduce food intake, thereby inducing weight loss(*43*). These appetite-inhibiting drugs have short-term side effects including nausea, vomiting and diarrhea; and raise the risk of cardiovascular disease, osteoporosis and other disorders(*41, 44*). Unlike these appetite-inhibiting drugs, leptolin administration elevated EE and reduced weight loss, but did not affect food intake. The anti-obesity drugs, such as semaglutide and tirzepatide, may lead to a loss of the lean body mass (including muscle and bone)(*41, 45*). Exercise can improve metabolic status and lead to a healthier proportion of lean-to-total body mass(*46*). Thus, the “exerkine” leptolin may have the potential to improve metabolic status, reduce body weight and fat pad weight without reducing lean body mass. In the present study, we did not observe apparent toxicity in muscle, bone and other major organs of the leptolin-treated mice. Of note, leptolin gene knockout or overexpression, or leptolin administration for 2 months did not alter the motor activity of mice, reflecting that leptolin did not affect food-seeking behavior. Overall, leptolin may be a promising anti-obesity agent with safety, and leptolin action may recapitulate the benefits of exercise in regard to metabolic health.

Human and mouse leptolin protein were 77% identical, compared to 83% identity for leptin, 83% identity for adiponectin, 79% identity for FGF21 and 59% identity for resistin. Both human and mouse leptolin expression was inversely correlated with obesity. These certainly imply a conserved function of leptolin in human and mouse. Functions of adipokines, such as leptin, adiponectin, FGF21 and resistin, are mediated by specific receptor-ligand interactions(*5, 9, 47, 48*). The identity of leptolin receptor is still unknown. Further studies will be needed to comprehend the leptolin receptor mechanisms through which leptolin elevates EE and prevents obesity.

In conclusion, we identify leptolin as a novel anti-obesity adipokine. Leptolin treatment may represent a promising therapeutic strategy against obesity and related disorders. The clinical utility of leptolin needs to be further explored.

## Supporting information

sMovie1

## Acknowledgments

We thank Bruce M. Spiegelman (Department of Cancer Biology, Dana Farber Cancer Institute; Department of Cell Biology, Harvard Medical School) for helpful suggestion. We thank Xing Zeng (Department of Cancer Biology, Dana Farber Cancer Institute; Department of Cell Biology, Harvard Medical School), Zhengwei Xie (Peking University International Cancer Institute, Health Science Center, Peking University) and Jianyuan Luo (Department of Medical Genetics, Center for Medical Genetics, Peking University) for helpful discussion. We thank Wenjuan Wang and Jianwei Wang (Department of Anatomy, Histology and Embryology, Peking University) for technical support. We thank Xian Wang and Shigong Zhu (Department of Physiology and Pathophysiology, Peking University) and Min Ye (Peking University School of Pharmaceutical Sciences) for kindly providing access to necessary equipment. We thank Rui Wei (Department of Endocrinology and Metabolism, State Key Laboratory of Female Fertility Promotion, Peking University Third Hospital, Beijing) for kindly providing plasma samples. We also thank the teams of the Novogene, the Advanced Cell Diagnostics, the SinoBiological and the Viewsolid Biotech for support of necessary techniques.

## Funding

National Natural Science Foundation of China (No. 81471064; R.Z.)

National Natural Science Foundation of China (No. 81670779; R.Z.)

National Natural Science Foundation of China (No. 81870590; R.Z.)

National Natural Science Foundation of China (No. 82170864; R.Z.)

National Key Research and Development Program of China (2017YFC1700402; R.Z.)

Beijing Municipal Natural Science Foundation (No. 7162097; R.Z.)

Beijing Municipal Natural Science Foundation (No. H2018206641; R.Z.)

Peking University Research Foundation (No. BMU20140366; R.Z.)

Scientific Project of Beijing Life Science Academy (No. 2023300CB0100; R.Z.).

## Author contributions

JL performed the experiment, analyzed the data, made the figures, did the literature search and wrote the paper. BW, ZS, XH, MH, YZ, YH, DL, WZ, LQ, KW, YL, YY, SY, XH, and TY participated in the study. RZ conceived the study, designed experiments, and wrote and edited the paper. All authors contributed to the study, and reviewed and approved the manuscript for submission. All authors accept full responsibility for the content of this paper.

## Data and materials availability

All data are available in the manuscript or the supplementary materials.

## Supplementary Materials

Materials and Methods

Fig. S1 to S22

Table S1

Movie S1

