## supplementary materials for "Identification of leptolin as a novel anti-obesity adipokine"

Science
AAAS

### Supplementary Materials for

#### Title: Identification of leptolin as a novel anti-obesity adipokine

Jiarui Liu[1], Bingwei Wang[1], Zhijie Su[1], Xiaoxv Han[2], Miao He[3], Yun Zhao[1], Yujia Hou[1], Daotong Li[1], Weiguang Zhang[1], Lihua Qin[1], Ke Wang[1], Yanchun Li[4], Yi Yan[5], Siwang Yu[6], Xiaoshuai Huang[3], Tairan Yuwen[2], Ruimao Zheng[1,7,8,9]*.

**The PDF file includes:**

Materials and Methods
Figs. S1 to S22
Table S1

**Other Supplementary Materials for this manuscript include the following:**

Movie S1

#### Materials and Methods

##### *Mice and Animal Care*

Mice were housed at 22 ± 1 °C with a 12-h light/dark cycle. Standard mouse chow and water were freely available except where otherwise indicated. All procedures were approved by the Institutional Care and Use Committee of the Peking University Health Science Center (approval number: LA2019340). All animals were sex- and age-matched, and littermates were used, as indicated in the figures. Animals were allocated to their experimental group according to their genotypes. Leptolin$^{+/+}$ and leptolin$^{-/-}$ mice were generated by crossing heterozygous leptolin$^{+/-}$ mice. Wild type (WT) and leptolin gene overexpression (leptolin-TG) mice were generated by crossing the C57BL/6n mice and leptolin-TG mice. C57BL/6n male mice were obtained from the Department of Laboratory Animal Science of Peking University Health Science Center. For diet-induced obesity studies, mice were placed on the 60 kcal% HFD (D12492i; Research Diets) at the age of 8 weeks. For thermoneutral studies, mice were housed at 30°C in a light-controlled climatic chamber. During all procedures of experiments, the number of animals and their suffering by treatments were minimized.

##### *Human Subjects*

These nonobese/obese human subjects were obtained from the Peking University Third Hospital (approval number: 202-226-02). Informed consent was obtained from all patients or their parents/ guardians. All data were kept confidential and processed anonymously. The research encompassed 55 samples of human blood plasma, procured from 27 obese individuals (BMI ≥ 28 kg/m²) and 28 nonobese individuals (18 < BMI < 28 kg/m²), with matching age and sex. Weight was measured before blood collection. The nonathletes/athletes human subjects were obtained from the Beijing Sport University (approval number: 2018018H). Informed consent was obtained from all participants. All data were kept confidential and processed anonymously. The research encompassed 32 samples of human blood plasma, procured from 16 athletes and 16 nonathletes with matching age and sex. All of the participants displayed apparent health, without any record of excessive alcohol consumption.

##### *Establishment of fat burning models*

Running exercise was carried out as previously described; Briefly, 10-week-old mice were housed individually with ad libitum food and water; One week before the experiment, mice were adapted on a motor treadmill at a speed of 10 m/min for 10 min on the first day and increased the exercise time by 10 min each day until it reached 60 min per day; Then, this exercise strength for mice was maintained for another two weeks. Swimming exercise was carried out as described; Briefly, 10-week-old mice were trained to swim for 10 min in warm (30 °C) water and twice a day; The swimming time were increased by 10 min each day until reaching 90 min per day; Then, this exercise strength for mice was maintained for another two weeks; We closely checked the mice to avoid anoxia and kept the water temperature at 30 ℃ to avoid hypothermia; After each test session, mice were towel-dried and placed back in their home cage. Cold exposure was conducted according to a previously established program; In brief, 10-week-old mice were housed individually with ad libitum food and water; Mice were placed at 4 ℃ for seven days before sacrificing.

##### *Whole Transcriptome Sequencing and Analysis*

Total RNA was extracted from iWAT of Ctrl, Run, Swim or Cold group mice using TRIzol reagent (TransGen Biotech). The quality of the RNA was determined with the NanoDrop 5500 (Thermo Fisher Scientific). For library preparation, 3 μg of total RNA/sample was used. Sequencing libraries were generated with NEBNext Ultra RNA Library Prep Kit for Illumina (New England Biolabs, Ipswich, MA). RNA molecules were selected using poly-T oligo-attached magnetic beads, fragmented, and reverse transcribed with the Elute, Prime, Fragment Mix. Then, end repair, A-tailing, adaptor ligation, and library enrichment were performed according to the manufacturer's instructions. RNA libraries were assessed for quality using the Agilent 2100 Bioanalyzer. The clustering of the index-coded samples was performed on a cBot Cluster Generation System using TruSeq PE Cluster Kit v3-cBot-HS (Illumina). Then, the RNA libraries were sequenced as 100-bp/50-bp paired-end runs on an Illumina HiSeq 2000/2500 platform. Differential expression analysis of two conditions/groups (two biological replicates per condition) was performed using the DESeq R package (1.10.1). The resulting P values were adjusted using the Benjamini-Hochberg approach for controlling the false discovery rate. Genes with an adjusted P value <0.05 found by DESeq were assigned as differentially expressed. Gene Ontology (GO) enrichment analysis of differentially expressed genes (DEGs) was performed using the GOseq R package, in which gene length bias was corrected. Samples were measured in Novogene Bioinformatics Technology Co., Ltd.

***Generation of leptolin-KO mice***

For generation of leptolin$^{-/-}$ (leptolin-KO) mice, Cas9/gRNAs were microinjected into the fertilized eggs. Injected eggs were implanted into surrogate C57BL/6n mothers to obtain offspring. Genotyping was performed to identify those harboring the tmem52 deficiency. Heterozygous mice were used to generate homozygous leptolin$^{+/+}$ and leptolin$^{-/-}$ mice. Primers for genotyping are as follows: leptolin-WT forward: 5′- AGA AGC AGG CAT TCC AGG TTG AG -3′, reverse: 5′- AGA GGC AGC AGC AGC AGG AT -3′, leptolin-KO forward: 5′- AGC AGG CAT TCC AGG TTG AG -3′, reverse: 5′- CAG TCC AGG TCT GTG GTC AC -3′.

***Generation of leptolin-TG mice***

For generation of pCAG-leptolin-P2A-EGFP transgenic mice, a vector encoding leptolin-P2A-EGFP under the control of cytomegalovirus (CMV) enhancer fused to the chicken beta-actin promoter (CAG promoter) was obtained. The plasmid was microinjected into the pronucleus of fertilized eggs. Injected eggs were implanted into surrogate C57BL/6 n mothers to obtain offspring. Genotyping was performed to identify those carrying the leptolin transgene. Protein levels of exogenous leptolin in plasma was approximately doubled in leptolin-TG mice (fig2d). Primers for genotyping are as follows: leptolin-TG forward: 5′- GCC ATA CTC CTG ATG CTT TTG T -3′, reverse: 5′- AGT TCA CCT TGA TGC CGT TCT -3′.

***Leptolin protein injection production***

Mouse tmem52 cDNA was cloned and subsequently sub-cloned into a pET-22B vector for expression in Escherichia coli. The fusion protein that was expressed in E. coli is 135 amino acids long comprising of a 6-amino-acid His tag on the N terminus and a 129-amino-acid leptolin fragment (68-196 amino acids). The His-leptolin were isolated from E. coli and allowed to bind to Ni-NTA His-Bind column. After extensive washing of the column in order to remove contaminating proteins, His-leptolin were eluted from the column using a 150-mM imidazole buffer. Then the His tag was cut, and the leptolin protein was further purified using size-exclusion columns and polymyxin B-based endotoxin-depletion columns (Detoxi-Gel Endotoxin Removing Gel by Thermo Scientific), to bring the final endotoxin concentration equal to or below 2 EU/ml. The buffer was exchanged into a PBS buffer.

***Administration of leptolin***

Administration of leptolin was carried out by i.p. injection. For i.p. treatment, mice received 100 μl of vehicle (PBS) for 4 days as acclimation before leptolin treatment. Then, leptolin was dissolved in PBS (100 μl; 0.005, 0.05, 0.5, and 5 mg/kg) and administered to mice once a day for 2 months; Vehicle groups received 100 μl of PBS during the course of the experiments. All treatments were performed within 90 min of the dark cycle.

***Real-Time Quantitative PCR***

RNA was extracted from the tissues using TransZol Up reagent (ET111-01, TransGen Biotech), followed by reverse transcription using a reverse transcription kit (AT311-03, TransGen Biotech). cDNAs were processed for real-time PCR using SYBR Green mix (AQ141-04, TransGen Biotech) with specific primers on a StepOnePlus real-time PCR System (Roche). All data were normalized with GAPDH.

***RNAscope***

Inguinal WATs were collected from mice, frozen in OCT compound (Sakura Finetek), and immediately followed by preservation at -80°C according to standard procedure. Tissue sections (SuperFrost Plus microslides) of 10-25 μm thickness were taken on a cryostat. The sections were immediately fixed in 4% paraformaldehyde solution for 15 minutes. Then, the sections were incubated sequentially with 50% ethyl alcohol for 5 minutes, 70% ethyl alcohol for 5 minutes, and 100% ethyl alcohol two times for 5 minutes and allowed to air dry on slides for 5 minutes. Sections were incubated with RNAscope® Hydrogen Peroxide for 2 hours at 40°C, and were washed in 1X Wash Buffer two times for 2 minutes at room temperature (RT). Sections were incubated with RNAscope® 2.5 AMP 1 for 30 minutes at 40°C, with RNAscope® 2.5 AMP 2 for 15 minutes at 40°C, with RNAscope® 2.5 AMP 3 for 30 minutes at 40°C, with RNAscope® 2.5 AMP 4 for 15 minutes at 40°C, with RNAscope® 2.5 Amp 5-RED for 30 minutes at RT, and with RNAscope® 2.5 Amp 6-RED for 15 minutes at RT. Then sections were incubated with RNAscope® RED working solution for 10 minutes at RT, and were washed in distilled water two times. Sections were incubated with 50% Hematoxylin staining solution for 2 minutes at RT, and then were incubated with 0.02% Ammonia water. Sections were mounted with VectaMont medium and analyzed on a microscope (Histology Facility of the Department of Anatomy, Histology and Embryology, Peking University).

***Total Protein Extraction and Western Blotting***

Proteins were extracted from iWAT, eWAT, BAT or blood plasma using a RIPA lysis buffer containing 0.5% NP-40, 0.1% sodium deoxycholate, 150 mm NaCl, 50 mm Tris–HCl (pH 7.4), phosphatase inhibitors (B15002, Bimake), and protease inhibitor cocktail (B14002, Bimake). Following 5 min of homogenization, lysates were centrifuged at 11 250 r for 15 min at 4 °C. Supernatants from the fat pads were used as protein extracts. The concentration of each sample was calculated by the BCA method and an equal amount of protein from each sample was added an equal amount of protein loading buffer, this buffer should contain 5% β-mercaptoethanol (vol/vol) or 20mmol/L TCEP (for western blotting of leptolin), and denatured by boiling at

100 °C for 5 min. Equal amounts of proteins were separated by 10% SDS-PAGE, transferred to NC membranes. The membranes were blocked for 2 h in 5% skim milk. The membranes were then incubated with the primary antibody in 5% BSA-TSBT at 4 °C. After overnight incubation, the membrane was washed three times in TBST for 15 min, followed by incubation with a secondary antibody in TBST with 5% skim milk for 2 h at room temperature. Following three cycles of 15 min washes with 1× TBST, the membranes were developed using a chemiluminescence assay. Intensities of the protein bands were quantified by ImageJ software. The antibodies used in this study include Anti-Pgc1α antibody (Bioss, bs-1832R), Anti-pHSL$^{(S660)}$ antibody (Bioss, bs-3358R), Anti-HSL antibody (Bioss, bs-0455R), Anti-α/β-Tubulin antibody (Cell Signaling Technology, 2148S), Anti-Leptolin antibody (designed by us and produced by SinoBiological), Anti-Rabbit IgG(H+L) (Biodragon, BF03008), and Anti-Mouse IgG(H+L) (Biodragon, BF03001).

***Immunofluorescence***

Inguinal WATs were collected from mice and immediately fixed in 4% paraformaldehyde solution for 48 h. Then, the samples were incubated sequentially with 20% sucrose and 30% sucrose in PBS for 2 d and frozen in OCT compound (Sakura Finetek). Tissue sections of 10-25 μm thickness were taken on a cryostat and allowed to air dry on slides, followed by processing or preservation at −80°C according to standard procedure. 3T3L1 cells were placed and cultured in confocal dishes, and fixed in 4% paraformaldehyde solution for 30 minutes. Frozen sections of tissues or confocal dishes of cells were subjected to leptolin or VAMP2 staining. Sections or dishes were washed in PBS for 10 min, followed by incubation in blocking solution (10% normal goat serum, 0.2% Triton X-100, 2% BSA, and PBS) for at least 1 h at room temperature. Primary antibodies were applied in blocking solution and incubated overnight at 4°C. Sections or dishes were washed at least three times with 5-min incubations in PBS plus 0.2% Triton X-100. Then, an Alexa Fluor 488–conjugated secondary antibody (1:200) or Alexa Fluor 594–conjugated secondary antibody (1:200) was applied in blocking solution and incubated at room temperature for 2 h, followed by five washes with PBS plus 0.2% Triton X-100, and nuclei were stained with DAPI. Sections or dishes were mounted with VECTASHIELD medium (Vector Laboratories) and analyzed on a microscope (Histology Facility of the Department of Anatomy, Histology and Embryology, Peking University). The antibodies used in this study include Anti-Leptolin antibody (designed by us and produced by SinoBiological), Anti-VAMP2 antibody (Proteintech, 67822-1-lg), Anti-Rabbit IgG/Alexa Fluor 488 (Abcam, ab150108), and Anti-Mouse IgG/Alexa Fluor 594 (Abcam, ab150108).

***Hematoxylin and Eosin Staining***

Animals were sacrificed and fats were immediately dissected and fixed in 4% paraformaldehyde solution for 48 h followed by cryopreservation in 25% sucrose solution (wt/vol) overnight and subsequent freezing in OCT compound (Tissue-Tek). Samples were stored in optimal cutting temperature compound (OCT) for frozen. Samples were sectioned, and H&E stained. The cell size was calculated by Image J.

***Metabolic Chamber***

Indirect calorimetry recording was performed using an indirect open-circuit calorimeter Oxylet Physiocage System (LE1305 Physiocage 00, LE405 O2/CO2 Analyzer, and LE400 Air Supply and Swithching; Panlab, Cornellà, Spain). Room air flowed through each chamber at a rate of 450 mL/min. The mice were placed in metabolic chambers with fresh food and water provided every day to acclimate for 24 hr. Before dark cycle, the authors started to monitor the oxygen consumption (VO2), carbon dioxide production (VCO2), respiratory-exchange-ratio (RER), energy expenditure (EE), and motor activity for 24 h. The O2 and CO2 levels were measured during 3-min sampling periods every 30 min, and data were analyzed with METABOLISM software (v2.2.01). Locomotor activity were measured using a two-dimensional infrared light beam. The VO2 and VCO2 were expressed in milliliters per minute per kilogram. The respiratory exchange ratio (RER) was determined by the ratio VCO2/VO2. The EE was calculated with the Weir equation. These metabolic parameters were adjusted for mouse adiposity. The mean values for RER, EE and motor activity of the dark cycle and light cycle were compared for each group.

***Insulin and Glucose Tolerance Tests***

For insulin and glucose tolerance tests (ITT and GTT, respectively), mice were fasted for 4 and 16 h, respectively. Mice were injected intraperitoneally with bovine insulin (Sigma, 0.5 units/kg) or with 20% glucose (2.0 g/kg). Blood samples were taken at 0, 30, 60, and 120min.

***Plasma parameter assays***

Whole blood was collected by cardiac puncture and transferred to ice-cold EP tubes. The tubes were centrifuged at 2,000g for 30 min at 4°C and stored at −80°C. The serum was used for western blot and measurement of insulin, TG and CHO levels. Insulin, triglycerides and cholesterol reagent kits were purchased from Biosino Bio-Technology and Science Inc.

***Body composition analysis***

Body composition analysis was performed using EchoMRI device (Echo Medical Systems, Houston, TX) to assess total body fat and lean mass of the mice.

***Cell maintenance and preparation***

293T cells were cultured in high-glucose DMEM (GIBCO, 11965092) supplemented with 10% fetal bovine serum (FBS) (GIBCO) and 1% 100 mM sodium pyruvate solution (Sigma-Aldrich, S8636) in an incubator at 37 °C with 5% CO2. 293T cells were transfected with a leptolin-EGFP vector. Transfections were performed using Lipofectamine$^{TM}$ 2000 (Thermo Fisher Scientific, 11668019) according to the manufacturer's instructions. Images or videos of cells were obtained using SR fluorescence-assisted diffraction computational tomography 24–36 h after transfection. 3T3L1 cells were cultured in high-glucose DMEM (GIBCO, 11965092) supplemented with 10% fetal bovine serum (FBS) (GIBCO) and 1% 100 mM sodium pyruvate solution (Sigma-Aldrich, S8636) in an incubator at 37 °C with 5% CO2. For preparation of differentiated 3T3L1 cells, 3T3L1 cells were cultured in high-glucose DMEM (GIBCO, 11965092) supplemented with 10% fetal bovine serum (FBS) (GIBCO), 0.5mM IBMX, 1μM insulin and 0.5μM dexamethasone.

***Bioinformatic Analysis of Human Datasets***

A total of 4 Ctrl/Exercise human sWAT datasets (GSE43471, GSE115645, GSE116801, GSE159809) and 2 large sample human sWAT datasets with BMI information (GSE135134, GSE70353) were identified, log2-transformation was applied as needed. In order to control for the effects of known and hidden covariates in each of the datasets, we used R/sva package to first adjust expression data for known factor covariates with the combat function and then estimate surrogates for hidden covariates with the sva function. Within each dataset, the expression values were further standardized to $\mu = 0$ and $\sigma = 1$. Scatter plots of BMI, and expression levels of tmem52 and genes associated with mitochondrial function and fat mobilization were displayed, and correlation analysis was performed. A total of 16 Normal weight/Obese human sWAT datasets (GSE141432, GSE166047, GSE165932, GSE152991, GSE159924, GSE24883, GSE44000, GSE55200, GSE59034, GSE88837, GSE156906, GSE48964, GSE205668, GSE29718, GSE133099, GSE94752) were identified, log2-transformation was applied as needed, and meta-analysis of tmem52 expression levels in these datasets was performed.

***Statistical Analysis***

Where indicated, data were expressed as mean ± standard error of means (SEM). Statistical significance was determined using Student t test (two-tailed) or one-way ANOVA. All data were analyzed using the appropriate statistical analysis methods with SPSS software (Windows version 26, IBM Analytics) or GraphPad Prism (Windows version 8.0, GraphPad Software). Significance was accepted at $^{*}P < 0.05$, $^{**}P < 0.01$, or $^{***}P < 0.001$. Sample sizes (n), statistical tests and p values are indicated in each figure legend.

Fig. S1.

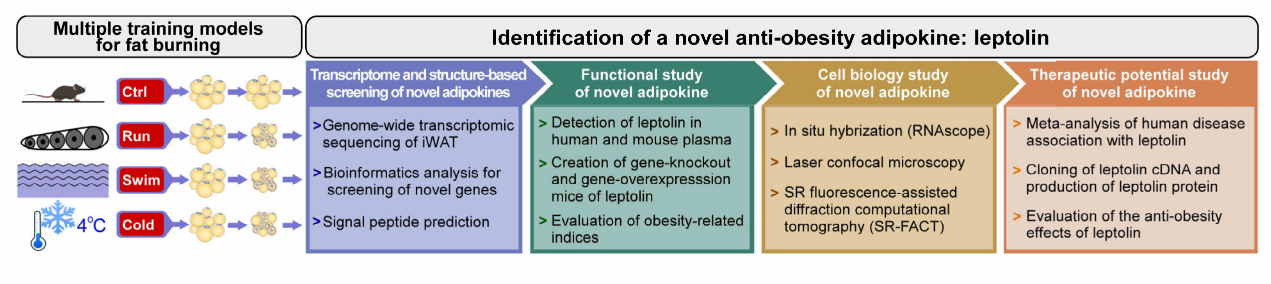


**Fig. S1. Schematic illustration shows the experimental strategy and workflow for identifying novel anti-obesity adipokines.**

**Fig. S2.**

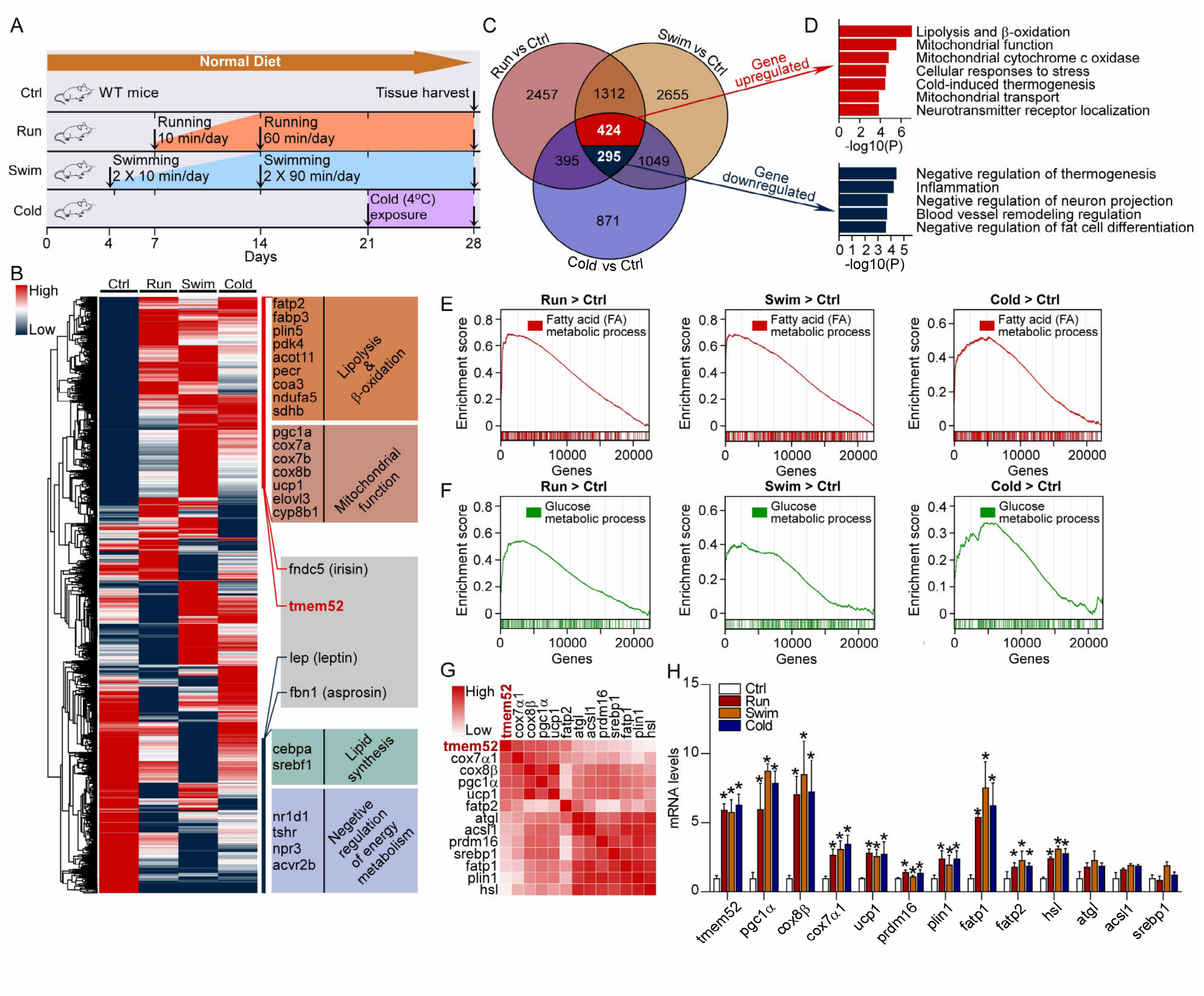


**Fig. S2. Tmem52 and lipolysis-related genes are activated under fat-burning conditions. (A)** Schematic illustration of experiments to establish fat-burning mouse models. **(B)** Hierarchical clustered heatmap of gene expression profiles in iWAT of ctrl, run, swim and cold groups; n = 3 per group. **(C)** Venn diagram of overlapping differential expression genes (DEGs). **(D)** Gene Ontology (GO) enrichment analysis based on the overlapping DEGs. **(E-F)** GSEA analysis based on the gene expression profiles in iWAT of ctrl, run, swim and cold groups. **(G)** Heatmap showing correlation between tmem52 and genes associated with mitochondrial function and fat mobilization. **(H)** Relative mRNA expression of the indicated genes in iWAT; n = 6 per group; Data were mean ± s.e.m.; One-way ANOVA with Bonferroni test was used for statistical analysis; *$P < 0.05$.

**Fig. S3.**

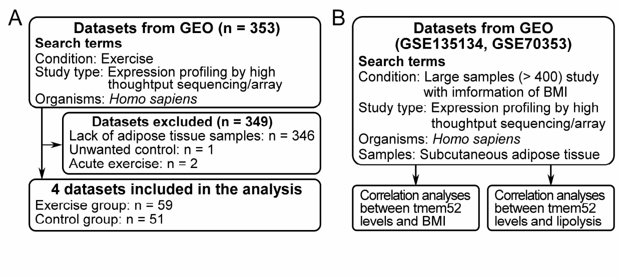


**Fig. S3. Screening workflow of human sWAT datasets. (A)** Schematic illustration of screening workflow of Ctrl/Exercise human sWAT datasets, 4 datasets were included in the analysis. **(B)** Schematic illustration of screening workflow of large sample human sWAT datasets with information of BMI, 2 datasets were included in the analysis.

**Fig. S4.**

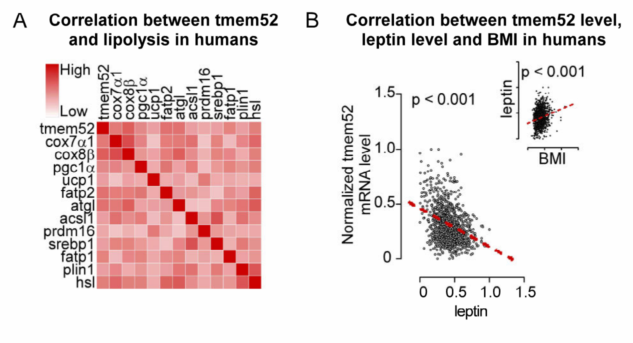


**Fig. S4. Tmem52 expression is positively correlated with lipolysis-promoting genes and negatively correlated with leptin in human sWAT. (A)** Heatmap showing correlation between tmem52 and genes associated with mitochondrial function and fat mobilization, based on the differently normalized large sample human datasets (n = 1203). **(B)** Scatter plots of BMI, and the mRNA levels of tmem52 and leptin in sWAT of differently normalized large sample human datasets (n = 1203); The red line showed correlation.

**Fig. S5.**

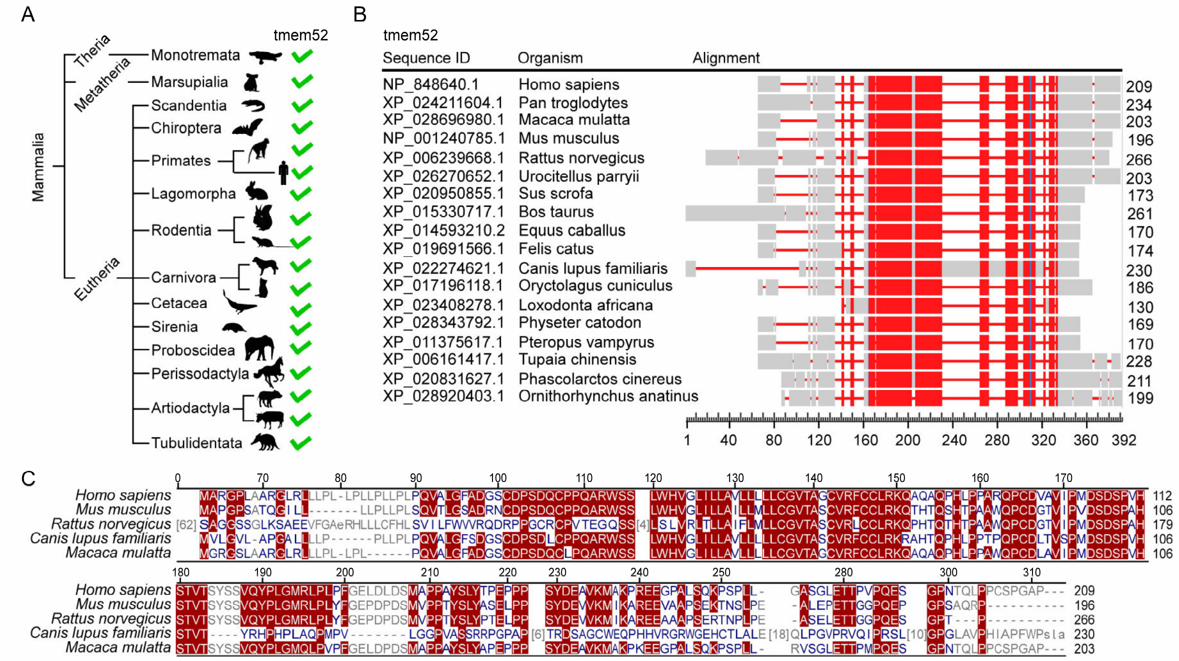


**Fig. S5. Unequivocal homologs and conserved amino acid sequence of tmem52 are identified in mammals. (A)** Phylogenetic tree analysis showing tmem52 gene in mammal species. **(B)** Multiple alignment of tmem52 of the mammal species. **(C)** Multiple alignment of tmem52 of the 5 species (*Homo sapiens*, *Mus musculus*, *Rattus norvegicus*, *Canis lupus familiaris*, *Macaca mulatta*).

#### Fig. S6.

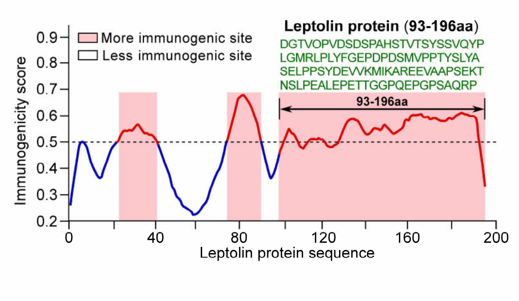


**Fig. S6. An immunogenic antigen is selected for the generation of a functional and highly specific anti-leptolin antibody.**

**Fig. S7.**

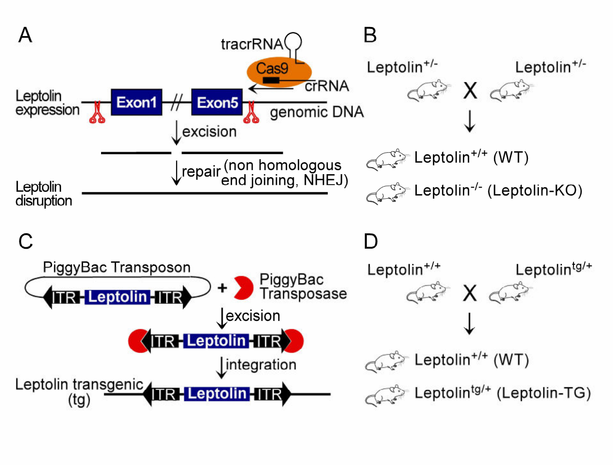


**Fig. S7. Generation of the leptolin-KO and leptolin-TG mice. (A)** Schematic of the knockout strategy for tmem52 gene. **(B)** Breeding strategy for generation of leptolin$^{+/+}$ mice (WT) and leptolin$^{-/-}$ mice (leptolin-KO). **(C)**Schematic of the overexpression strategy for tmem52 gene. **(D)** Breeding strategy for generation of wild type mice (WT) and leptolin gene overexpression mice (leptolin-TG).

**Fig. S8.**

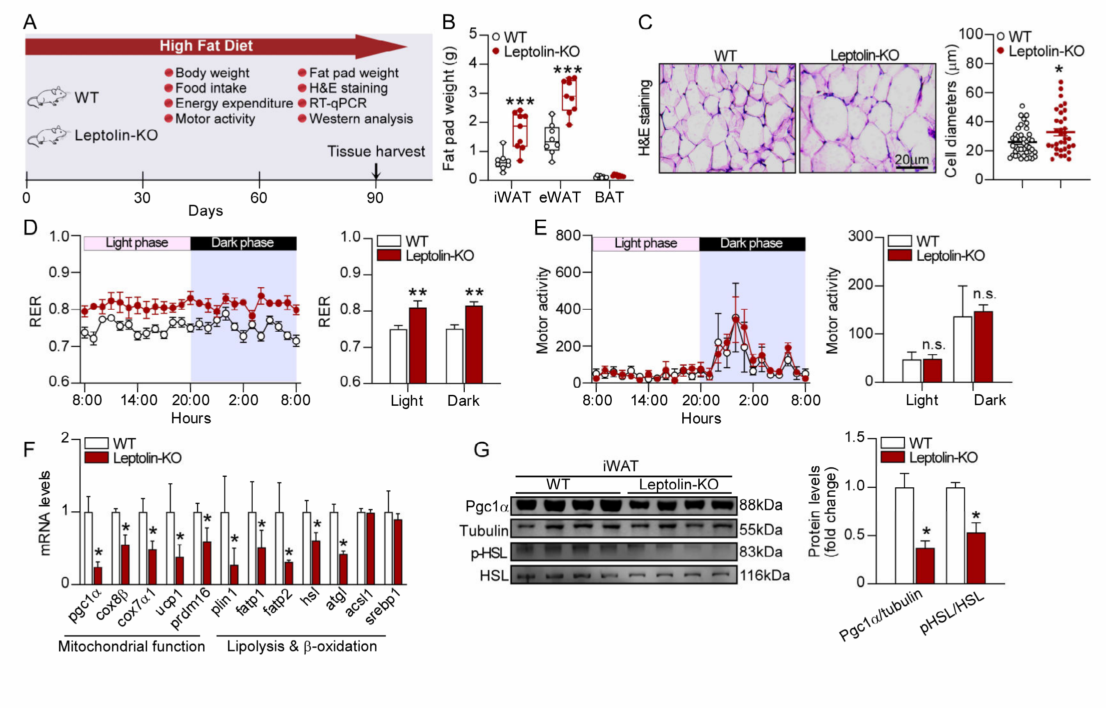


**Fig. S8. Leptolin deficiency prevents lipolysis and reduces EE. (A)** Schematic illustration of experiments. **(B)** Fat pad weight; WT, n = 8, Leptolin-KO, n = 9. **(C)** Representative images of hematoxylin and eosin (H&E) staining of iWAT and the size profiling of adipocytes from iWAT; Scale bar indicated 20 µm. **(D-E)** RER **(D)** and motor activity **(E)**; n = 5 per group. **(F)** Relative mRNA expression (fold change) of genes associated with mitochondrial function and fat mobilization in iWAT; n = 6 per group. **(G)** Representative immunoblots of pgc1α, tubulin, p-HSL$^{(S660)}$ and HSL from iWAT, and the quantified ratio of pgc1α/tubulin and p-HSL$^{(S660)}$/HSL; n = 8 per group. Data were mean ± s.e.m.; Student's t-test was used for statistical analysis; *p < 0.05, **p < 0.01, ***p < 0.001, n.s., not significant.

**Fig. S9.**

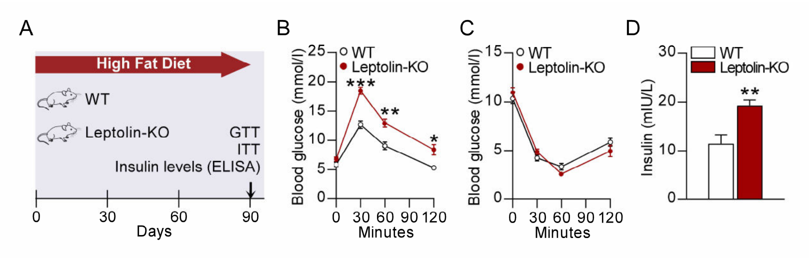


**Fig. S9. Leptolin deficiency dampens glucose tolerance in mice. (A)** Schematic illustration of experiments. **(B-C)** Glucose tolerance test **(B)** and insulin tolerance test **(C)**; WT, n = 8, Leptolin-KO, n = 9. **(D)** Plasma insulin level; WT, n = 6, Leptolin-KO, n = 8. Data were mean ± s.e.m.; Student's t-test was used for statistical analysis; $*p < 0.05$, $**p < 0.01$, $***p < 0.001$.

**Fig. S10.**

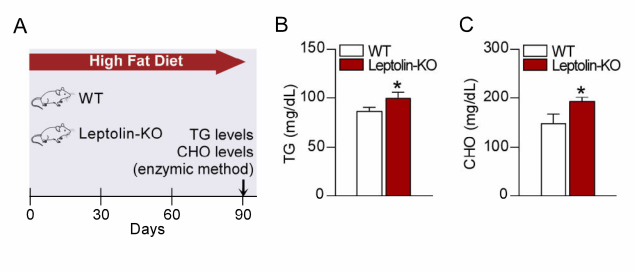


**Fig. S10. Leptolin deficiency elevates the triglyceride and total cholesterol levels in plasma. (A)** Schematic illustration of experiments. **(B)** Plasma triglyceride levels; n = 5 per group. **(C)** Plasma total cholesterol levels; WT, n = 6, Leptolin-KO, n = 7. Data were mean ± s.e.m.; Student's t-test was used for statistical analysis; *$p < 0.05$.

**Fig. S11.**

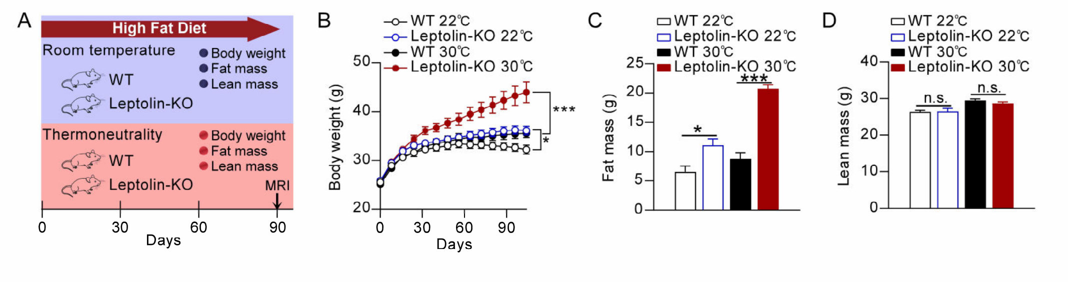


**Fig. S11. Leptolin deficiency increases the susceptibility to HFD-induced obesity in mice at thermoneutrality. (A-D)** 8-week-old WT and leptolin-KO mice were housed at either 22°C or 30°C and fed a high-fat-diet. **(A)**Schematic illustration of experiments. **(B)** Body weight; WT 22°C, n = 8, Leptolin-KO 22°C, n = 9, WT 30°C, n = 9, Leptolin-KO 30°C, n = 9. **(C-D)** Fat mass **(C)** and lean mass **(D)**; n = 5 per group. Data were mean ± s.e.m.; One-way ANOVAs were used for statistical analysis followed by Bonferroni's post hoc test; *p < 0.05, ***p < 0.001, n.s., not significant.

**Fig. S12.**

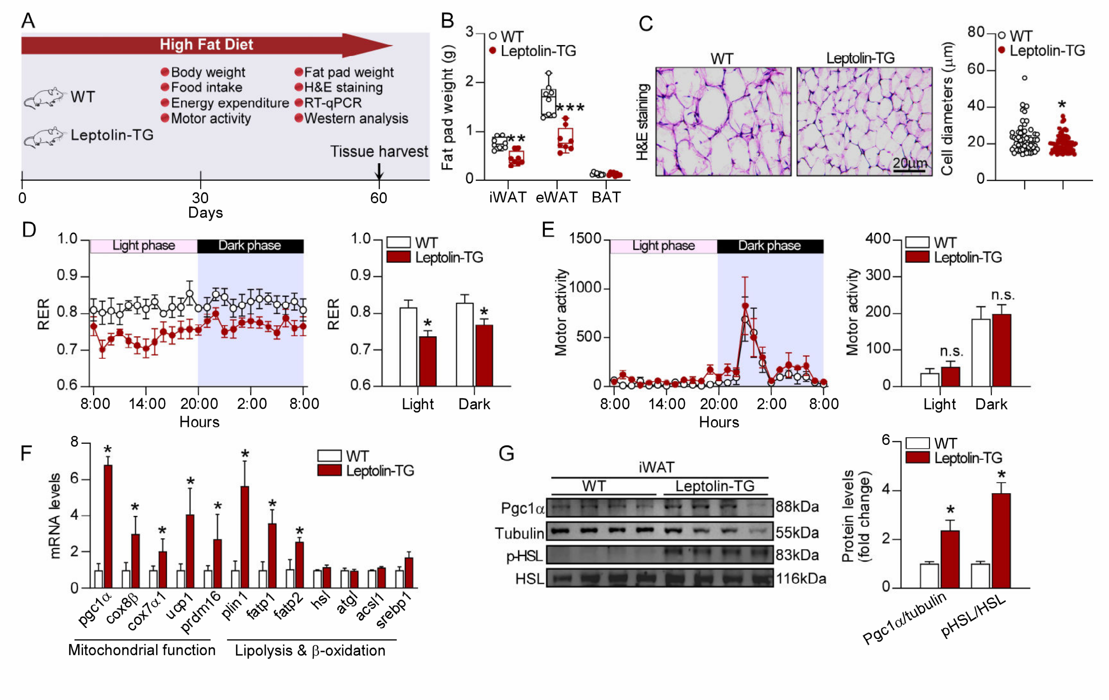


**Fig. S12. Leptolin gene overexpression promotes lipolysis and elevates EE. (A)** Schematic illustration of experiments. **(B)** Fat pad weight; WT, n = 8, Leptolin-TG, n = 9. **(C)** Representative images of hematoxylin and eosin (H&E) staining of iWAT and the size profiling of adipocytes from iWAT; Scale bar indicated 20 μm. **(D-E)** RER **(D)** and motor activity **(E)**; n = 5 per group. **(F)** Relative mRNA expression (fold change) of genes associated with mitochondrial function and fat mobilization in iWAT; n = 6 per group. **(G)** Representative immunoblots of pgc1α, tubulin, p-HSL$^{(S660)}$ and HSL from iWAT, and the quantified ratio of pgc1α/tubulin and p-HSL$^{(S660)}$/HSL; n = 8 per group. Data were mean ± s.e.m.; Student's t-test was used for statistical analysis; *p < 0.05, **p < 0.01, ***p < 0.001, n.s., not significant.

**Fig. S13.**

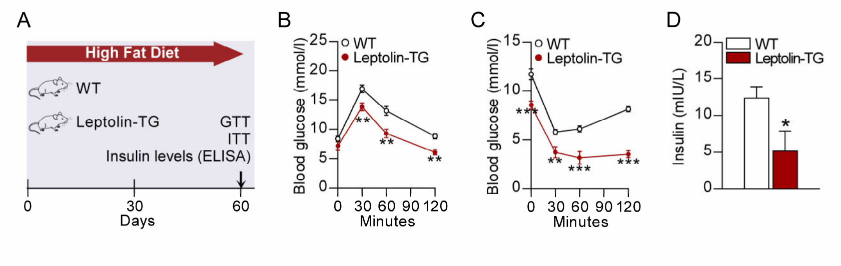


**Fig. S13. Leptolin gene overexpression improves glucose tolerance and insulin sensitivity in mice. (A)** Schematic illustration of experiments. **(B-C)** Glucose tolerance test **(B)** and insulin tolerance test **(C)**; WT, n = 6, Leptolin-TG, n = 6. **(D)** Plasma insulin level; WT, n = 5, Leptolin-TG, n = 5. Data were mean ± s.e.m.; Student's t-test was used for statistical analysis; *$p < 0.05$, **$p < 0.01$, ***$p < 0.001$.

**Fig. S14.**

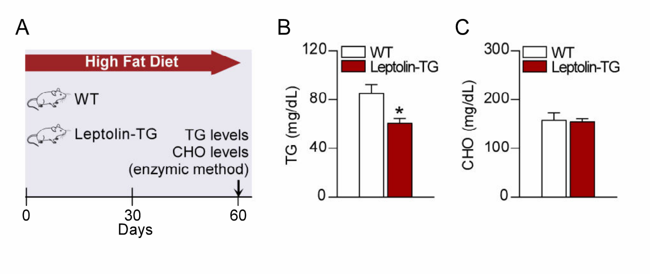


**Fig. S14. Leptolin gene overexpression reduces the triglyceride levels in plasma.** **(A)** Schematic illustration of experiments. **(B)** Plasma triglyceride levels; WT, n = 5, Leptolin-TG, n = 5. **(C)** Plasma total cholesterol levels; WT, n = 5, Leptolin-TG, n = 8. Data were mean ± s.e.m.; Student's t-test was used for statistical analysis; $^*p < 0.05$.

**Fig. S15.**

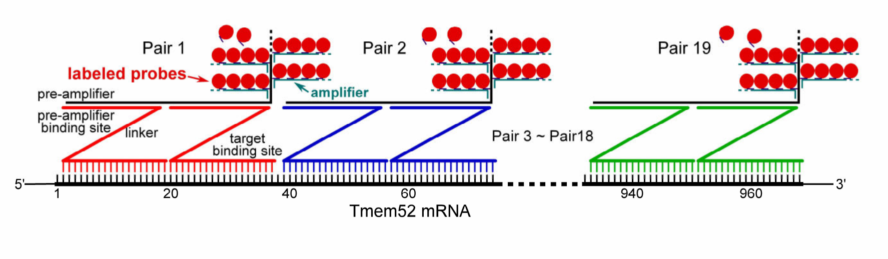


**Fig. S15. Schematic illustration shows the high sensitivity and specificity of RNAscope assay.**

Fig. S16.

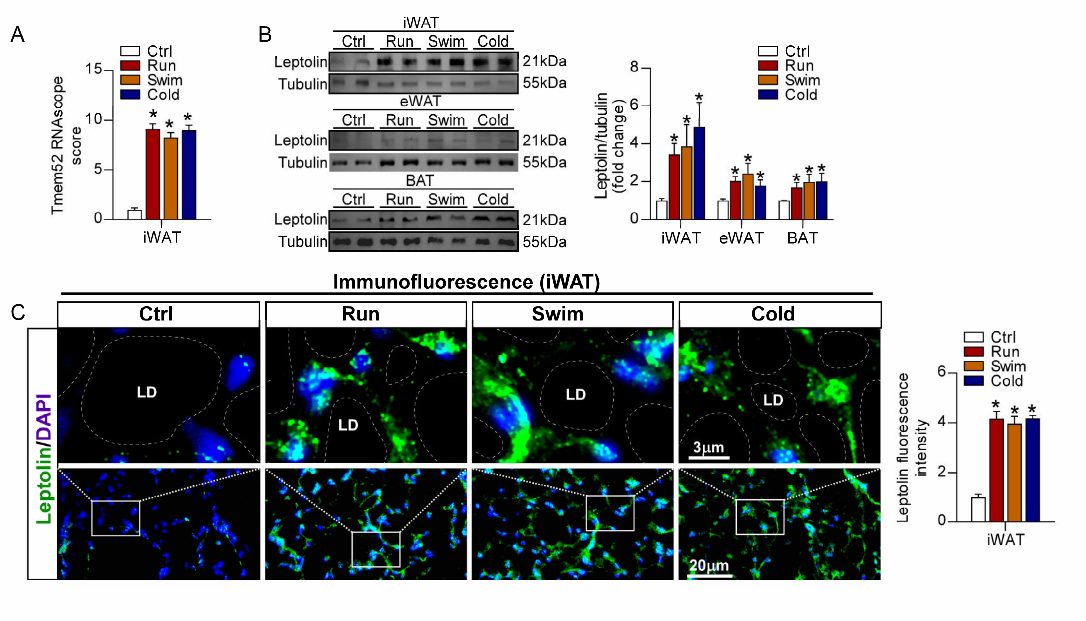


**Fig. S16. Leptolin is an adipose tissue-derived protein induced by fat-burning conditions. (A)** RNAscope score of tmem52 in iWAT of mice under fat-burning conditions. **(B)** Representative immunoblots of leptolin and tubulin from iWAT, eWAT and BAT, and the quantified ratio of leptolin/tubulin; n = 6 per group. **(C)** Representative immunofluorescence images of leptolin in iWAT, and the fluorescence intensity of leptolin; Scale bar, 3 μm (upper panel) and 20 μm (down panel). Data were mean ± s.e.m.; The one-way ANOVAs were used for statistical analysis followed by Bonferroni's post hoc test; *P < 0.05.

**Fig. S17.**

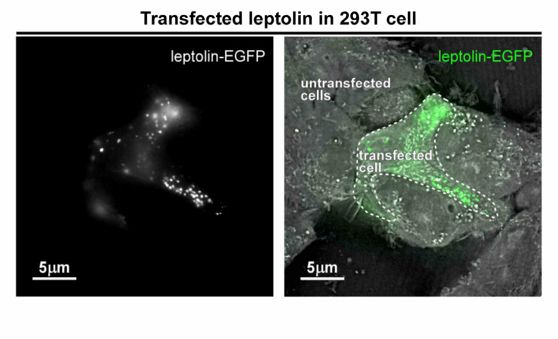


**Fig. S17. SR-FACT technique detects the leptolin-EGFP signals in vesicles of live 293T cells transfected with plasmid.**

**Fig. S18.**

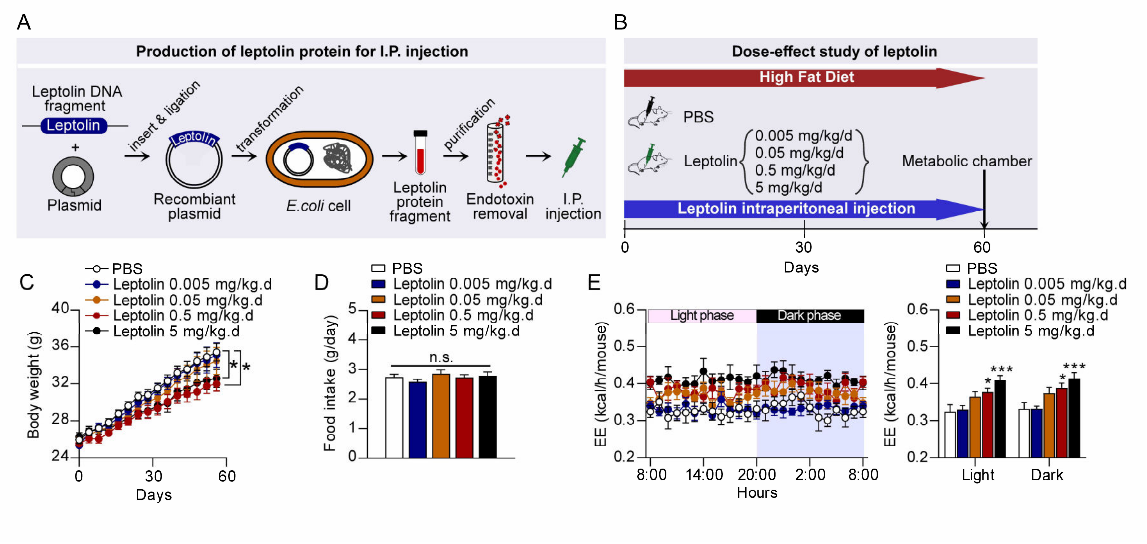


**Fig. S18. The dose-effect study of leptolin I.P. injection in treating obesity. (A)** Schematic of production strategy for leptolin protein injection. **(B)** Schematic illustration of experiments: 8-week-old mice were fed a high-fat-diet, received PBS for 4 days as acclimation, then were intraperitoneally injected with PBS or leptolin at dosages of 0.005-5 mg/kg/d daily for 2 months; PBS, n = 8, Leptolin 0.005 mg/kg/d, n = 9, Leptolin 0.05 mg/kg/d, n = 9, Leptolin 0.5 mg/kg/d, n = 8, Leptolin 5 mg/kg/d, n = 8. **(C)** Body weight. **(D)** Daily food intake. **(E)** Indirect calorimetry was performed to quantify EE during complete 24 hr light-dark cycles; n = 6 per group. Data were mean ± s.e.m.; One-way ANOVAs were used for statistical analysis followed by Tukey's post hoc test; *p < 0.05, ***p < 0.001, n.s., not significant.

Fig. S19.

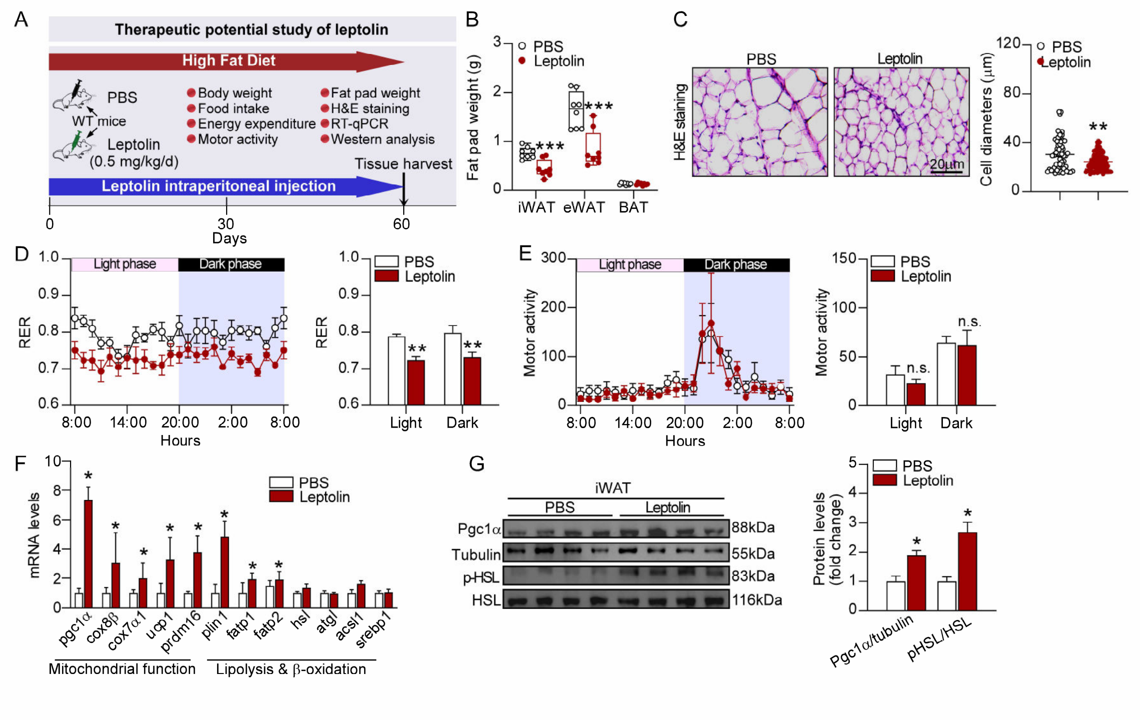


**Fig. S19. Leptolin administration promotes lipolysis and elevates EE. (A)** Schematic illustration of experiments. **(B)** Fat pad weight; PBS, n = 8, Leptolin, n = 8. **(C)** Representative images of hematoxylin and eosin (H&E) staining of iWAT and the size profiling of adipocytes from iWAT; Scale bar indicated 20 µm. **(D-E)** RER **(D)** and motor activity **(E)**; n = 6 per group. **(F)** Relative mRNA expression (fold change) of genes associated with mitochondrial function and fat mobilization in iWAT; n = 6 per group. **(G)** Representative immunoblots of pgc1α, tubulin, p-HSL$^{(S660)}$ and HSL from iWAT, and the quantified ratio of pgc1α/tubulin and p-HSL$^{(S660)}$/HSL; n = 8 per group. Data were mean ± s.e.m.; Student's t-test was used for statistical analysis; *$p < 0.05$, **$p < 0.01$, ***$p < 0.001$, n.s., not significant.

**Fig. S20.**

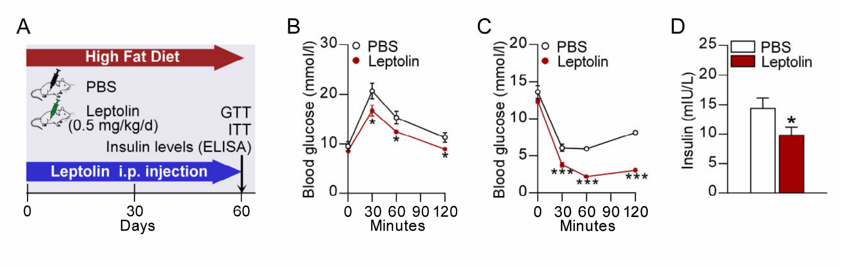


**Fig. S20. Leptolin administration improves glucose tolerance and insulin sensitivity in mice. (A)** Schematic illustration of experiments. **(B-C)** Glucose tolerance test **(B)** and insulin tolerance test **(C)**; n = 8 per group. **(D)** Plasma insulin level; PBS, n = 6, Leptolin, n = 8. Data were mean ± s.e.m.; Student's t-test was used for statistical analysis; *p < 0.05, ***p < 0.001.

**Fig. S21.**

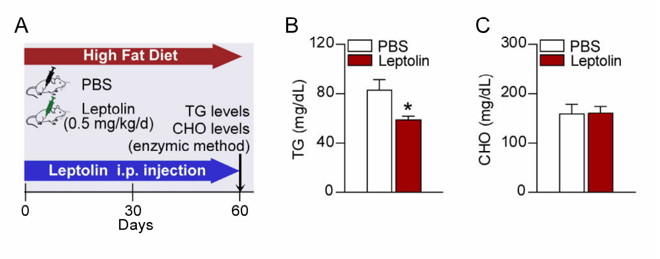


**Fig. S21. Leptolin administration reduces the triglyceride levels in plasma. (A)** Schematic illustration of experiments. **(B)** Plasma triglyceride levels; n = 6 per group. **(C)** Plasma total cholesterol levels; PBS, n = 5, Leptolin, n = 7. Data were mean ± s.e.m.; Student's t-test was used for statistical analysis; *$p < 0.05$.

#### Fig. S22.

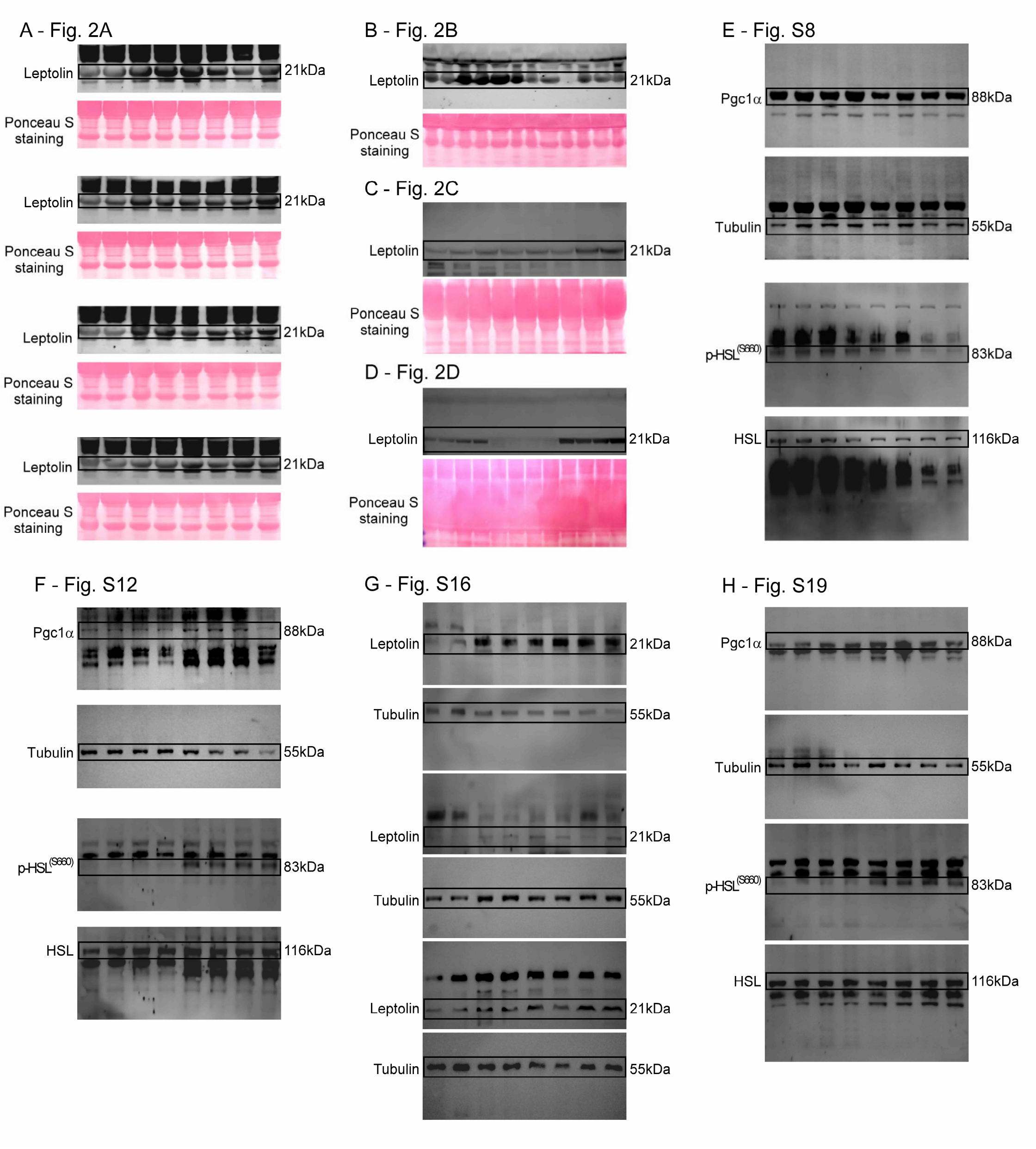


**Fig. S22. Original full western blot images.**

**Table S1. Primers used in the present study**

| Primer | Forward Primer 5'-3' | Reverse Primer 5'-3' |
|---|---|---|
| *tmem52* | TGCTCGCCATACTCCTGATG | CTGTGTGCGGGGCTATCAC |
| *pgc1α* | AGCCGTGACCACTGACAACGAG | GCTGCATGGTTCTGAGTGCTAAG |
| *cox8β* | GAACCATGAAGCCAACGACT | GCGAAGTTCACAGTGGTTCC |
| *cox7α1* | CAGCGTCATGGTCAGTCTGT | AGAAAACCGTGTGGCAGAGA |
| *ucp1* | ACTGCCACACCTCCAGTCATT | CTTTGCCTCACTCAGGATTGG |
| *prdm16* | CAGCACGGTGAAGCCATTC | GCGTGCATCCGCTTGTG |
| *plin1* | GGGACCTGTGAGTGCTTCC | GTATTGAAGAGCCGGGATCTTTT |
| *fatp1* | CGCTTTCTGCGTATCGTCTG | GATGCACGGGATCGTGTCT |
| *fatp2* | TCCTCCAAGATGTGCGGTACT | TAGGTGAGCGTCTCGTCTCG |
| *hsl* | GATTTACGCACGATGACACAGT | ACCTGCAAAGACATTAGACAGC |
| *atgl* | GGATGGCGGCATTTCAGACA | CAAAGGGTTGGGTTGGTTCAG |
| *acsl1* | TGCCAGAGCTGATTGACATTC | GGCATACCAGAAGGTGGTGAG |
| *srebp1* | TGACCCGGCTATTCCGTGA | CTGGGCTGAGCAATACAGTTC |

**Movie S1. SR-FACT technique detects the leptolin-EGFP signals in vesicles of live 293T cells transfected with plasmid.**
